# Hill-Based Reformulation of the Hodgkin-Huxley Model for Mapping Gating Changes to Neuronal Excitability

**DOI:** 10.64898/2026.06.06.730614

**Authors:** Batoul Saab, Jihad Fahs, Arij Daou

**Affiliations:** Neurophysiology and Computational Neuroscience Group, Biomedical Engineering Program, American University of Beirut, Lebanon; Department of Electrical and Computer Engineering, American University of Beirut, Lebanon; Harvard Radcliffe Institute for Advanced Study, Harvard University, Cambridge, USA

## Abstract

Conductance-based models link ion-channel gating to neuronal excitability, but mutation reports often provide steady-state voltage shifts without sufficient kinetic measurements to specify a unique model. We introduce a Hill-based reformulation of selected Hodgkin–Huxley (HH) gating relationships and evaluate its usefulness under this low-information constraint. The hybrid formulation retains canonical HH-like spike waveforms and frequency–current behavior after a modest remapping of applied current, while providing an explicit mapping from reported activation descriptors to model parameters under specified assumptions. An A863P kinetic-sensitivity analysis shows how steady-state constraints and kinetic assumptions jointly shape excitability predictions in both classical HH and Hill-based implementations. The Hill formulation makes this mapping reproducible through explicit parameter relations, whereas refitting classical HH rates requires additional fitting choices; both approaches still require kinetic assumptions. Retrospective benchmarks using the Nav1.7 mutations A863P, F1449V, and A1632G show that the Hill-based model captures the qualitative increase in excitability but overestimates the magnitude of threshold reduction. Classical HH comparisons for A863P demonstrate that similar qualitative predictions can also be obtained under specified kinetic assumptions. These findings support the Hill formulation as a reproducible framework for mapping reported gating changes onto HH-like excitability, without establishing superior quantitative prediction of mutation effects. Quantitative transfer to mutation-specific neuronal behavior requires additional kinetic and cellular constraints.

## Introduction

The Hodgkin-Huxley (HH) model remains one of the foundational mathematical frameworks in neuroscience for describing the ionic basis of action potential generation in excitable cells. Its conductance-based architecture continues to underlie modern single-compartment, multicompartment, and temperature-scaled neuronal models^1,2,2–6^. Adapting such models to a channel mutation is often a low-information problem: published measurements may report shifts in half-activation or half-inactivation voltage and slope, without providing the full voltage-dependent activation, inactivation, and recovery kinetics required for a unique dynamical model. The challenge is therefore to translate available measurements into explicit model perturbations while identifying the assumptions needed to complete that translation. Classical HH rate expressions can encode such changes, although their empirical constants do not always correspond directly to the reported steady-state quantities. A more explicit parameterization may facilitate mapping and reproducibility, but cannot supply kinetic information that was not measured.

Existing modeling strategies offer complementary tools. Reduced models such as Morris-Lecar and FitzHugh-Nagumo improve geometric and analytical tractability^7–10^, whereas phenomenological models such as Izhikevich reproduce broad firing repertoires^11,12^, but neither is designed to identify mutation-specific ionic kinetics from sparse voltage-clamp data. Boltzmann-type parameterizations retain a conductance-based interpretation and are widely used in voltage-clamp fitting and multicompartment modeling^13–18^. Their midpoint and slope parameters can be related directly to experimental measurements, providing a clear interpretation of gating changes. Classical HH rate functions can also be shifted or refitted, while steady-state Boltzmann curves can be combined with independently specified time constants. A fair comparison must distinguish the fit to a voltage-dependent relationship, the rule used to map experimental shifts, and the assumptions used to determine kinetics. Improved fitting or simpler coordinates alone do not establish more accurate prediction of neuronal excitability.

We investigate a Hill-based family of sigmoidal substitutions as an alternative coordinate system for this mapping. Hill functions describe saturating biological input–output relationships in diverse settings^19–25^. The Hill parameterization describes each selected sigmoidal relationship using a midpoint parameter and a Hill coefficient that controls its steepness. Here, the Hill coefficient is a phenomenologically steepness parameter, rather than evidence of molecular cooperativity. We distinguish explicit mapping of steady-state activation shifts from assumptions about inactivation kinetics: changing a selected inactivation rate does not independently prescribe the steady-state inactivation curve and its time constant. Both the Hill and classical HH formulations therefore require a stated kinetic rule when only steady-state measurements are available. We fit Hill functions to the HH-derived sigmoidal *m*_∞_, *n*_∞_ and *β*_ℎ_ relationships, benchmark their fits against Boltzmann, tanh, erf, and arctan alternatives, and test whether the resulting hybrid model preserves spike waveforms, frequency-current behavior, excitability regimes, and spike-features^1–5,23,26–33^. We use the Nav1.7-A863P channelopathy as a retrospective benchmark for translating experimentally reported sodium activation and inactivation shifts into both formulations^34^. We extend this retrospective comparison to F1449V and A1632G using limited reported gating information, and examine sensitivity to kinetic assumptions for A863P. These benchmarks test whether both formulations reproduce the direction of reported excitability changes and how closely they reproduce their magnitude; they are not prospective validation. Our hypothesis is that the Hill coordinates retain HH-like excitability and make the perturbation mapping easier to specify and reproduce. We do not assume that this mapping will outperform classical HH or Boltzmann-based alternatives quantitatively, particularly when mutation-specific kinetics and cellular context are incompletely constrained.

## Results

To evaluate how replacing classical HH rate expressions with Hill-type functions affects neuronal dynamics, we assessed the reformulation at four levels: derived gating kinetics, spike generation and input-output behavior, parameter-space organization of firing regimes and spike features, and retrospective benchmarks using the Nav1.7 mutations A863P, F1449V, and A1632G, including an A863P kinetic-sensitivity analysis. Together, these analyses ask whether the Hill-based model preserves the core behavior of the original HH system while providing a more transparent parameterization for tuning experimentally meaningful gating changes.

### 3.1 The Hill-based reformulation preserves steady-state gating while selectively reshaping channel time constants

To determine how the Hill-based reformulation alters the actual gating dynamics of the HH model, we compared the derived steady-state gating variables and time constants between the original and modified formulations (Figure 1). This analysis is important because the gating variables in the HH formalism do not evolve directly according to the raw *α_x_*(*V*) and *β_x_*(*V*) rate functions alone, but rather according to the relaxation form given by equation (3) in the Methods section.

**Figure 1.**
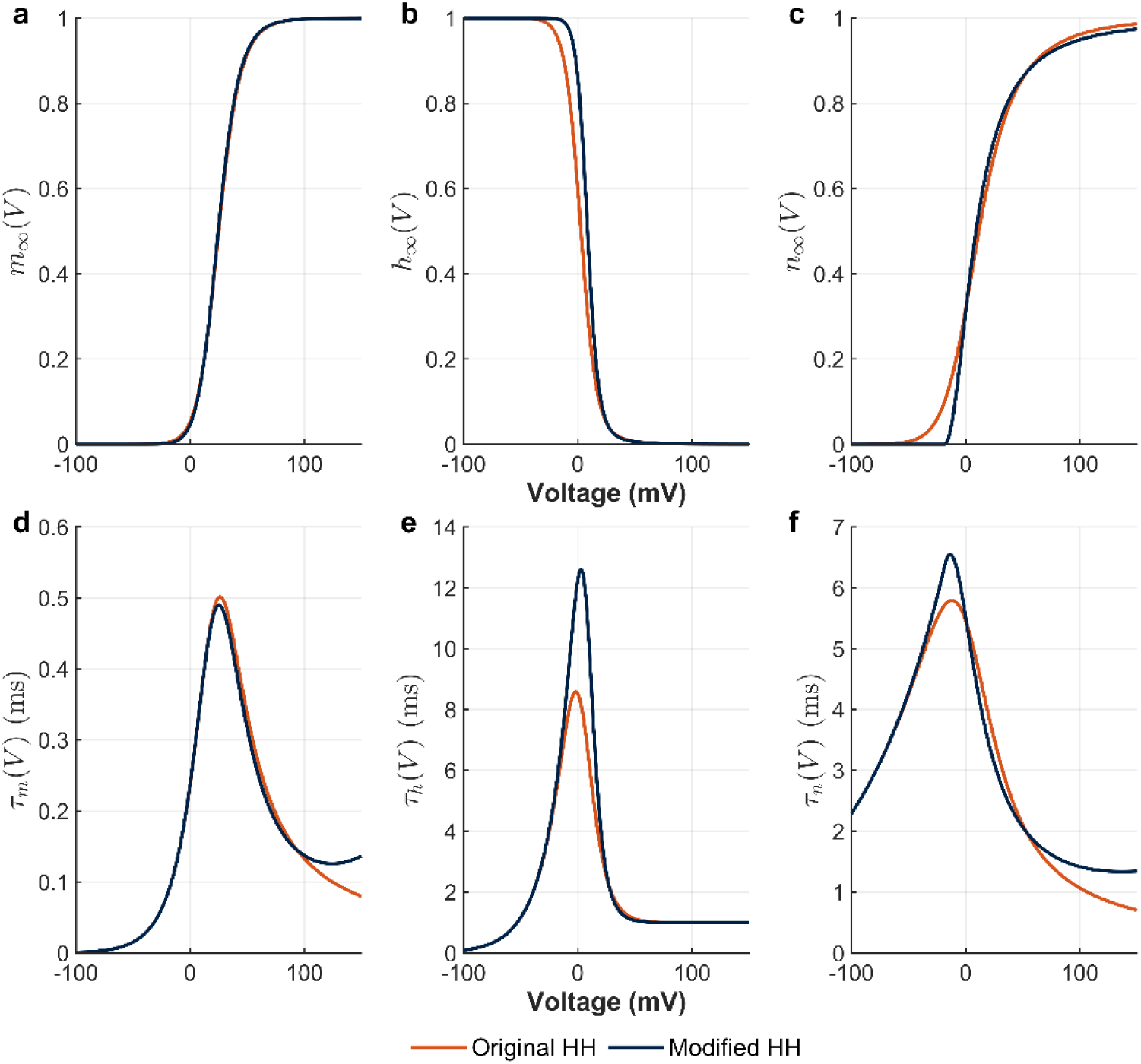
Derived gating consequences of the Hill-based kinetic reformulation. Comparison of the steady-state gating variables and time constants in the original Hodgkin-Huxley (HH) model (orange) and the modified Hill-based formulation (navy). Panels **a-c** show steady-state activation and inactivation curves m_∞_(V), ℎ_∞_(V), and n_∞_(V), respectively. Panels **d-f** show the corresponding time constants τ_m_(V), τ_ℎ_(V), and τ_n_(V). The modified model closely preserves the steady-state voltage dependence of all three gating variables, with only modest differences in the position and sharpness of the transition regions. By contrast, larger differences emerge in the time constants, particularly for τ_ℎ_(V) and τ_n_(V), indicating that the principal dynamical effect of the reformulation is to reshape the kinetics of relaxation more than the steady-state occupancy itself. These results provide a procedural link between the fitted Hill-based gating relationships and the preserved HH-like firing dynamics of the modified model.

Figures 1a-c show the steady-state activation and inactivation curves for the *m*, ℎ, and *n* gates, respectively. Across all three variables, the modified model remains close to the original HH formulation, indicating that the Hill-based reformulation largely preserves the voltage dependence of steady-state gating. In particular, *m*_∞_(*V*) is nearly superimposed between the two models, while ℎ_∞_(*V*) and *n*_∞_(*V*) show only modest differences in the position and sharpness of their transition regions. These results indicate that the reformulation leaves the steady-state occupancy structure of the classical HH model largely intact ^1,23,27,35^.

Figures 1d-f reveal more substantial differences in the time constants. Whereas *τ_m_*(*V*) remains comparatively similar between the two formulations, both *τ*_ℎ_(*V*) and *τ_n_*(*V*) are altered more appreciably in the modified model. In particular, the modified formulation produces a larger peak in *τ*_ℎ_(*V*) over the intermediate voltage range, indicating slower inactivation dynamics in this regime, and also reshapes *τ_n_*(*V*), with a higher peak and a slower voltage-dependent decay than in the original HH model. These effects indicate that the main dynamical consequence of the reformulation lies more in changing how rapidly the gates approach their steady-state values than in changing those values themselves.

Taken together, these results show that the Hill-based reformulation is conservative at the level of steady-state gating but more flexible at the level of relaxation kinetics. This distinction is important. It suggests that the modified model preserves the fundamental voltage dependence of activation and inactivation while selectively reshaping the temporal recruitment of sodium and potassium conductances. Such changes provide a plausible explanation for why the reformulated system retains canonical HH-like spike waveforms and firing patterns, yet exhibits a modest shift in effective excitability relative to the classical model ^2,12,36,37^.

### 3.2 Canonical Hodgkin-Huxley spiking dynamics are preserved after a modest remapping of effective input current

Having established that Hill functions provide the best compact sigmoidal description of the relevant HH-derived gating relationships, we next asked whether this reformulation preserves the dynamical behavior of the classical HH model at the level of excitability and input-output structure. Representative voltage traces comparing the original and modified models under matched firing conditions are provided in Supplementary Figure 1, where the modified model reproduces the spike timing and waveform morphology of the original HH system after a modest increase in applied current. The main-text analysis in Figure 2 therefore focuses on quantifying this correspondence across input levels rather than illustrating individual examples alone ^8,12,36,37^.

**Figure 2.**
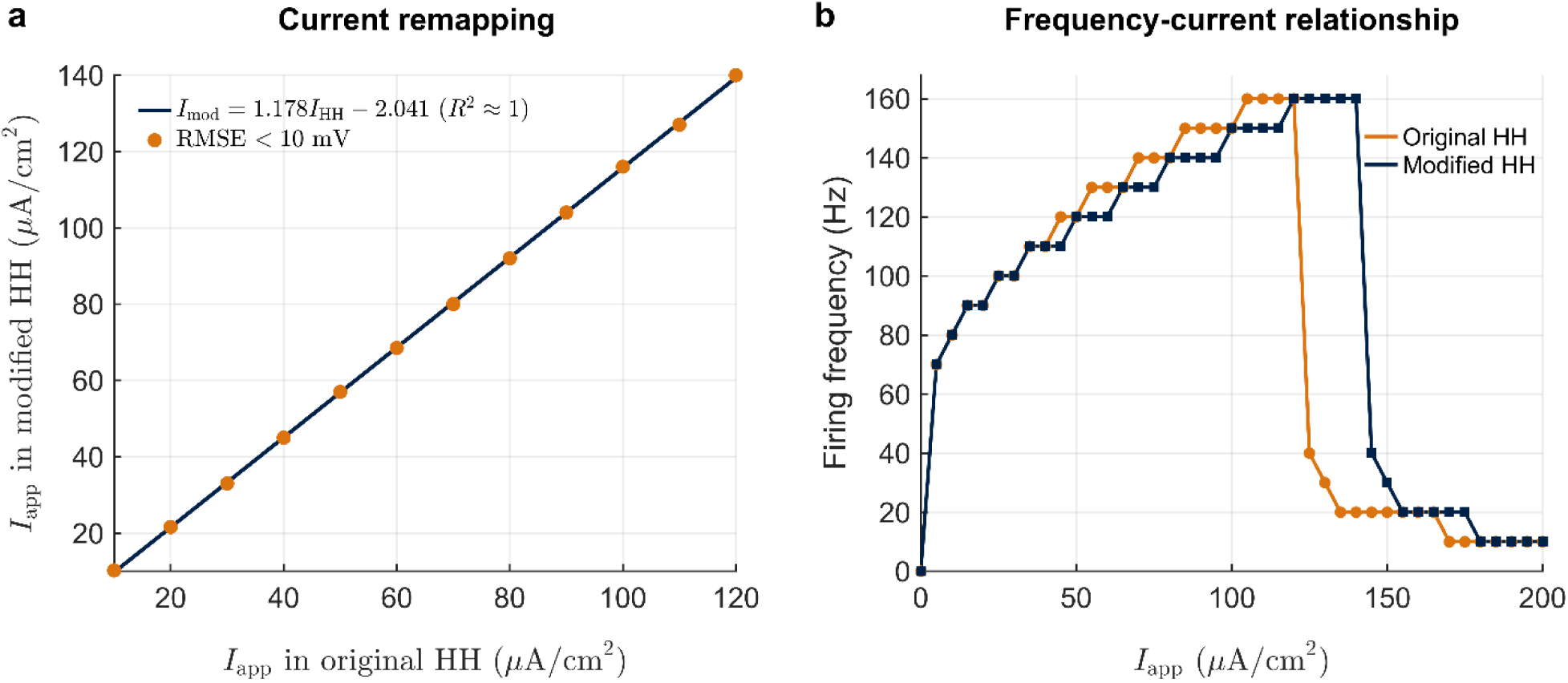
Quantitative preservation of canonical Hodgkin-Huxley excitability after Hill-based kinetic reformulation. **(a)** Mapping between the applied current in the original Hodgkin-Huxley (HH) model and the corresponding current in the modified model required to produce matched responses. Orange markers denote current pairs 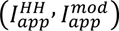 satisfying RMSE<10 mV, and the navy line shows best linear fit, 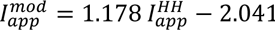, yielding a coefficient of determination almost equal to 1. The near-linear relationship indicates that the primary effect of the reformulation is a modest rescaling of excitability rather than a qualitative change in firing dynamics. **(b)** Frequency-current (f–I) relationships for the original and modified HH models under step-current stimulation. The modified model closely reproduces the global input–output behavior of the original model, including firing onset, the progressive increase in firing rate over intermediate inputs, the high-frequency plateau, and the breakdown of sustained repetitive firing at strong drive, with only a modest shift along the input axis. Together, these results show that the Hill-based reformulation preserves the overall dynamical organization of classical HH excitability while slightly altering its effective operating point. Representative matched voltage traces are shown in Supplementary Fig. 1.

We first quantified the change in effective drive required for the modified model to reproduce responses matched to those of the original HH model (Figure 2a). Across the explored range, the relationship between the applied current in the original model and the corresponding current in the modified model remained strikingly linear, with the best-fit mapping given by

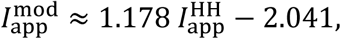

yielding a coefficient of determination close to 1. Matched response pairs were selected such that the root mean square error (RMSE) between the voltage traces of the two models remained below 10 mV, indicating close agreement in spike count and waveform properties. This near-affine mapping shows that the principal dynamical consequence of the Hill-based reformulation is not a qualitative distortion of firing behavior, but rather a smooth renormalization of excitability. Thus, the modified model preserves the broad input-output organization of the classical HH model while operating at a modestly shifted current scale.

We then asked whether this close correspondence extends beyond individual traces to the broader input-output behavior of the model. To this end, both the original and modified systems were stimulated with step currents ranging from 0 to 200 *μ*A/cm^2^ for 100 ms, and firing frequency was computed from spike counts over a fixed analysis window (Figure 2b). The resulting frequency-current (*f*– *I*) curves show that the modified model closely reproduces the global firing behavior of the original HH model, including firing onset, the progressive increase in discharge rate over intermediate current levels, a broad high-frequency plateau, and the abrupt loss of sustained high-rate firing at strong drive. The modified model shows a modest rightward shift in the current axis, fully consistent with the remapping quantified in Figure 2a, but preserves the overall gain structure and accessible firing-rate range of the original system ^8,12,36,37^.

This preservation is important because it indicates that replacing selected HH kinetic terms with Hill-type functions does not disrupt the canonical firing phenotype of the model. Instead, the reformulated system retains the broad structure of the frequency-current relationship while differing from the original formulation primarily through a modest and highly structured change in effective excitability. Thus, the gain in parameter interpretability introduced by the Hill-based representation is achieved without sacrificing the core electrophysiological behaviors captured by the classical HH framework.

### 3.3 Hill-parameter space is organized into structured regimes of spontaneous and stimulus-evoked excitability

To move beyond one-dimensional parameter sweeps and determine how each Hill parameter interacts with external drive to shape neuronal output, we constructed two-dimensional excitability maps in the (*I*_app_, *θ*) plane, where *θ* denotes one of the nine parameters of the modified model (Figure 3). For each panel, a single Hill parameter was varied across its admissible range while the applied current was swept over a broad interval, and the resulting activity was classified into five dynamical regimes: silent, single spike, tonic firing, spontaneous firing, and depolarization block. This analysis reveals the global structure of excitability in the modified model and shows how the Hill-based parameterization organizes neuronal behavior into contiguous and interpretable dynamical domains ^12,35,36,38^.

**Figure 3.**
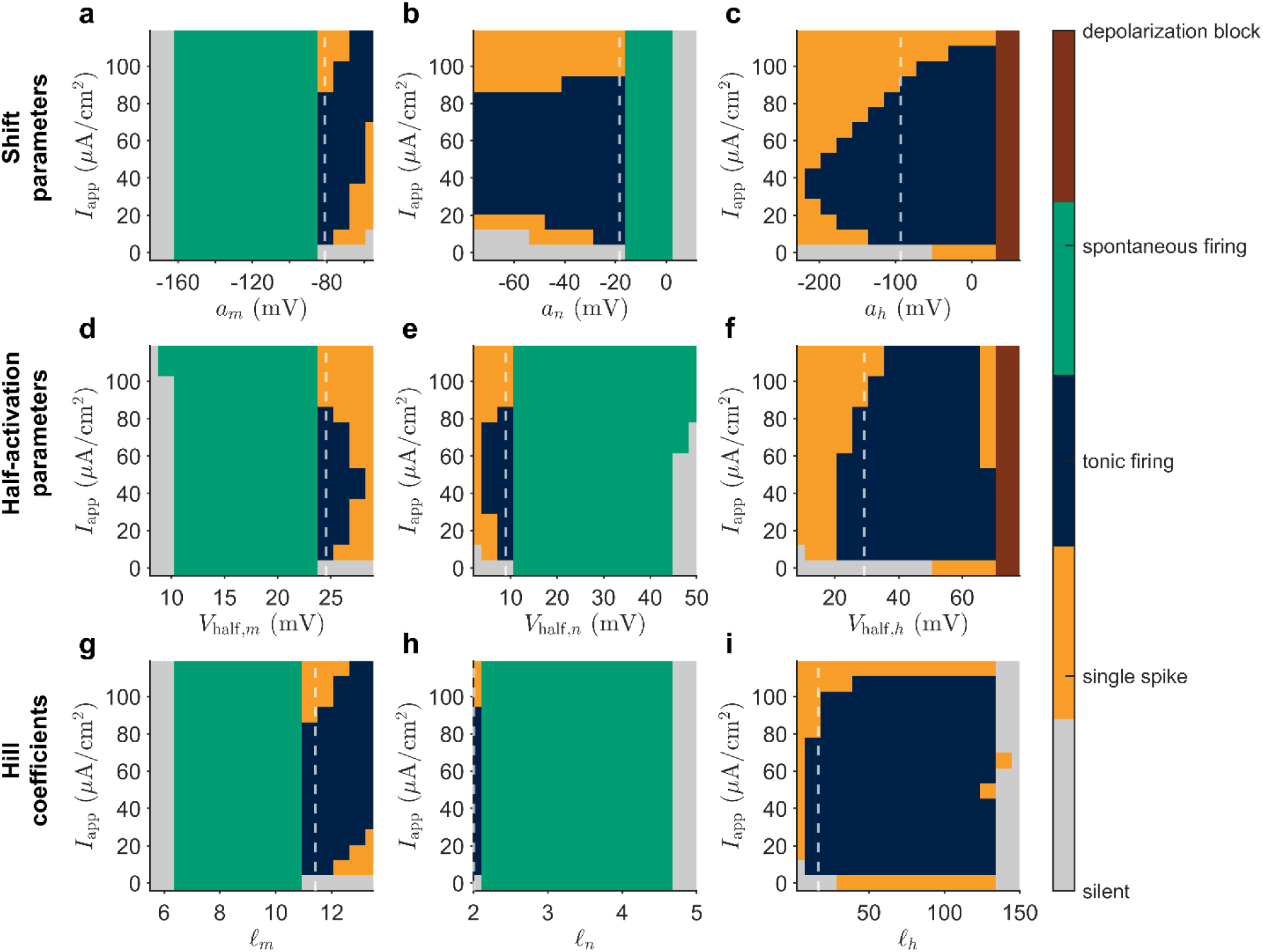
Excitability regime maps for the modified Hodgkin-Huxley model. Two-dimensional regime maps in the (*I*_app_, *θ*) plane for the nine Hill parameters of the modified model, with one parameter varied at a time and all others held fixed. Panels **a-c** show the shift parameters *a_m_*, *a_n_*, and *a*_ℎ_; panels **d-f** show the Hill midpoint parameters *V*_half,*m*_, *V*_half,*n*_, and *V*_half,ℎ_; and panels **g-i** show the Hill coefficients l*_m_*, l*_n_*, and l_ℎ_. Simulations were performed for 200 ms with a step current applied between 50 and 150 ms. Activity was classified as silent, single spike, tonic firing, spontaneous firing, or depolarization block. These maps show that the Hill parameters organize neuronal behavior into broad, contiguous dynamical regimes and locate transitions between silence, driven firing, autonomous activity, and firing failure.

Within the sampled parameter grid, Figures 3a-i show that regime boundaries are largely contiguous and orderly across parameter space. Rather than producing many isolated islands of activity or highly fragmented transitions, most parameters generate broad regions corresponding to distinct firing modes. These contiguous regions describe the sampled grid rather than establishing robustness to arbitrary parameter perturbations. Instead, the Hill parameters define a navigable excitability landscape in which gradual changes in a single functional quantity can move the model reproducibly between qualitatively different dynamical states ^30–33,39^.

The maps also reveal clear differences between the functional classes of Hill parameters. Sodium-activation parameters (Figures 3a, 3d, and 3g) show broad spontaneous-firing domains over one portion of parameter space and tonic-firing domains over another, separated by relatively sharp transitions. Altering the horizontal placement, half-activation voltage, or steepness of the sodium activation sigmoid therefore strongly affects whether the model remains rest-stable or enters self-sustained repetitive firing. Potassium activation parameters (Figures 3b, 3e, and 3h), particularly the half-activation and steepness terms, also exert strong control over the topology of the excitability map, in some cases generating very broad spontaneous-firing regions, indicating that the placement and steepness of potassium activation are highly influential determinants of resting-state stability under the modified formulation.

By contrast, the inactivation-related parameters (Figures 3c, 3f, and 3i) exhibit a qualitatively different organization. In these panels, broad regions of tonic firing occupy the center of parameter space, while spontaneous activity is largely absent or strongly reduced. Instead, one sees more pronounced transitions between silence, single-spike responses, tonic firing, and, at more extreme values, depolarization block. This suggests that the inactivation-related parameters act less as triggers of autonomous oscillation and more as regulators of the model’s ability to sustain repetitive firing under drive. The structured and localized appearance of depolarization block further supports the idea that the Hill parameters act as meaningful control axes for distinct forms of excitability failure rather than simply shifting spike features in an undifferentiated way ^1,40–42^. The depolarization block is most clearly associated with portions of the *a*_ℎ_ and *V*_half,ℎ_ maps, where the tonic-firing region terminates in a well-defined block regime at large parameter values and strong input. This is systematically sensible: altering sodium inactivation can profoundly affect the ability of the system to recover between spikes, and therefore determine whether repetitive firing is sustained or collapses into a depolarized non-spiking state ^41,43,44^.

The regime organization is consistent with the classical HH model’s Type II excitability, in which transition from silence to repetitive firing occurs via a subcritical Hopf bifurcation ^8,12,37,45,46^. Here, the Hill coordinates organize the observed activity changes along explicit parameter axes, although these maps alone do not establish the bifurcation structure of the modified model. Formal continuation analysis of Hopf and saddle-node loci remains a natural extension ^8,12,26,36–38^.

Taken together, these results show that the Hill-based reformulation does more than reproduce classical Hodgkin-Huxley-like firing under a new parameterization. It reorganizes the model’s dynamical behavior into structured regime maps defined by parameters with clear methodological meaning. Horizontal shifts, half-(in)activation voltages, and sigmoid steepnesses provide interpretable axes for moving between silence, single spiking, tonic driven firing, spontaneous activity, and depolarization block. This converts the kinetic parameter space into a more intelligible coordinate system for navigating neuronal dynamical regimes ^30,31,33,39^.

### 3.4 The Hill-based formulation yields an interpretable spike-feature landscape

To obtain a representative readout of repetitive firing, we compared spike features across positions in the train and found that the second spike most closely matched the mean of subsequent spikes across all four features - amplitude, threshold, width, and after-hyperpolarization (AHP) - in both models (Supplementary Figure 2). The first spike deviates systematically due to onset-specific transients that do not reflect sustained firing ^36,41,42,47^. We therefore use the second spike as a standardized proxy for early sustained discharge throughout the following analysis.

Figure 4 compares heat maps of these four second-spike properties for the original HH model and the modified formulation. For each sweep, one parameter was varied across its biologically admissible range while all others were held fixed at a reference configuration (black stars). Parameter ranges were selected to ensure biologically realistic repetitive firing - defined as non-damping spike trains with at least three spikes exceeding 0 mV - and are summarized in Supplementary Tables 5 and 6. All results were obtained under a constant applied current of 20 μA/*cm*^2^, and color scales are matched within each feature column across both models to permit direct comparison.

**Figure 4.**
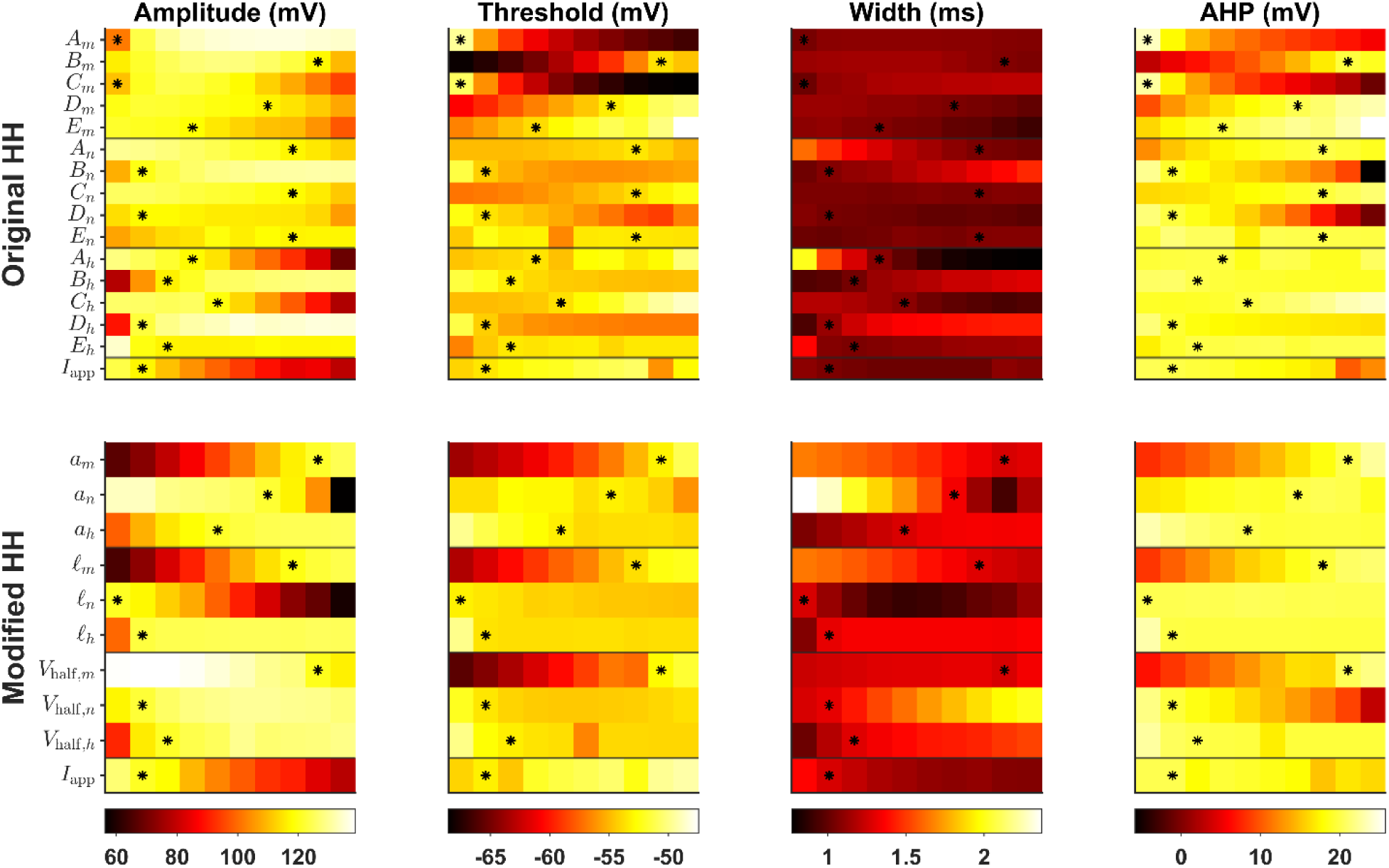
Parameter sensitivity of spike-waveform features in the original and Hill-based Hodgkin-Huxley models. Heat maps show how variation of individual model parameters affects four second-spike features: amplitude, threshold, width, and after-hyperpolarization (AHP). The upper block shows the original Hodgkin-Huxley (HH) model, and the lower block shows the modified Hill-based HH formulation. Within each block, each row corresponds to one varied parameter, while all remaining parameters were held fixed at their reference values. Black stars indicate the reference parameter value used for baseline simulations. The final row in each block shows the effect of varying the applied current. Color scales are matched within each feature column across the original and modified models, allowing direct comparison of the accessible feature ranges between parameterizations. The modified formulation preserves modulation of the major spike-waveform features while organizing this modulation through nine tunable Hill parameters corresponding to horizontal shift, Hill midpoint, and steepness.

A first key result is that the modified model preserves control over the same broad electrophysiological output dimensions classically associated with HH-type models, while doing so through nine tunable Hill parameters. In the original HH system, waveform tuning is distributed across fifteen empirical constants, *A* through *E* for the *m*, *n*, and ℎ gates, whose effects are often difficult to interpret functionally because each parameter enters indirectly through fitted rate laws. By contrast, the modified formulation reduces the kinetic description to nine tunable Hill parameters grouped by clear functional role: horizontal shifts (*a_m_*, *a_n_*, *a*_ℎ_), Hill midpoint voltages (*V*_1/2,*m*_, *V*_1/2,*n*_, *V*_1/2,ℎ_), and steepness coefficients (l*_m_*, l*_n_*, l_ℎ_). Figure 4 shows visually organized parameter-feature relationships along these coordinates. This comparison concerns the parameters varied in the sweeps; retained classical rates remain part of the hybrid model, and landscape smoothness was not formally quantified.

The clearest example is provided by spike amplitude and threshold, which in the modified model are strongly and systematically governed by parameters associated with sodium activation. In the amplitude maps, *a_m_*, *V*_1/2,*m*_, and l*_m_* generate broad and orderly gradients, indicating that shifting or steepening the sodium activation sigmoid produces predictable changes in the height of the action potential ^41,48^. The same logic extends to threshold, where the modified model reveals clearer gradients aligned with sodium-activation parameters and a more coherent dependence on current drive. In the original HH model, comparable effects are distributed across multiple empirical constants, producing correspondingly more diffuse and harder-to-interpret patterns. Related patterns are evident for spike width and AHP: the modified model spans a wider and more continuous range of widths, and AHP values vary more systematically across the parameter set, suggesting that the interplay between inward activation, delayed outward recruitment, and recovery is represented more transparently. Width and AHP provide additional waveform readouts, although these maps do not establish improved prediction of recordings from specific cell types ^43,49,50^.

A second major result is that the modified model accesses a broad spike-feature space while remaining compact. Within the admissible ranges explored here, the Hill-based formulation reaches lower spike amplitudes and wider spike widths than the original HH model, expanding the set of accessible waveform phenotypes without leaving regimes that satisfy the biological firing criteria. This comparison should be interpreted cautiously because the admissible ranges were defined independently for each model to preserve realistic repetitive firing within each respective formulation. Accordingly, the principal advantage claimed here is not a range-independent proof of superiority, but the more organized mapping between interpretable gating parameters and electrophysiological features. A formal normalized comparison of feature-space coverage and landscape smoothness remains a natural extension of the present simulation framework.

Taken together, Figure 4 shows that the Hill-based reformulation reorganizes the admissible excitability landscape into physiologically meaningful axes that exert clearer control over spike morphology. The principal advance is not only that HH-like dynamics are preserved, but that the mapping from kinetic parameters to electrophysiological phenotype can be inspected along the selected Hill coordinates, yielding a model that remains compact while providing a more interpretable description of waveform modulation.

### 3.5 A Nav1.7-A863P channelopathy benchmark tests biologically motivated gating shifts

To test whether the reformulation can encode a biologically characterized channel perturbation, we used the Nav1.7-A863P mutation associated with inherited erythromelalgia as a benchmark^34^. Harty et al. reported that A863P shifts Nav1.7 activation in the hyperpolarizing direction and steady-state fast inactivation in the depolarizing direction, increasing channel availability near rest and producing hyperexcitability in dorsal-root-ganglion (DRG) neurons^34^.

The benchmark was designed to test qualitative biological correspondence rather than to reconstruct the full DRG conductance repertoire. The experimentally reported gating shifts were imposed in both formulations. In the Hill-based model, the shifts were encoded directly through the sodium activation and inactivation-related Hill parameters. In the original HH consistency comparator, activation targets and the Hill-derived inactivation curve were reproduced by coordinated adjustment of the empirical rate parameters while holding complementary retained rates fixed. Activation and inactivation perturbations were first tested separately and then together.

Both implementations in this consistency comparison reproduced the expected direction of the A863P phenotype (Supplementary Figure 3; Figure 5b). In the original HH comparator, the A863P/WT rheobase ratio was 0.450 for the activation shift alone, 0.926 for the inactivation shift alone, and 0.397 for the combined perturbation. The modified model gave closely matched values of 0.460, 0.917, and 0.407, respectively. Thus, the hyperpolarizing activation shift accounted for most of the reduced threshold in this simplified benchmark, whereas the depolarizing inactivation shifts alone produced a smaller effect.

**Figure 5.**
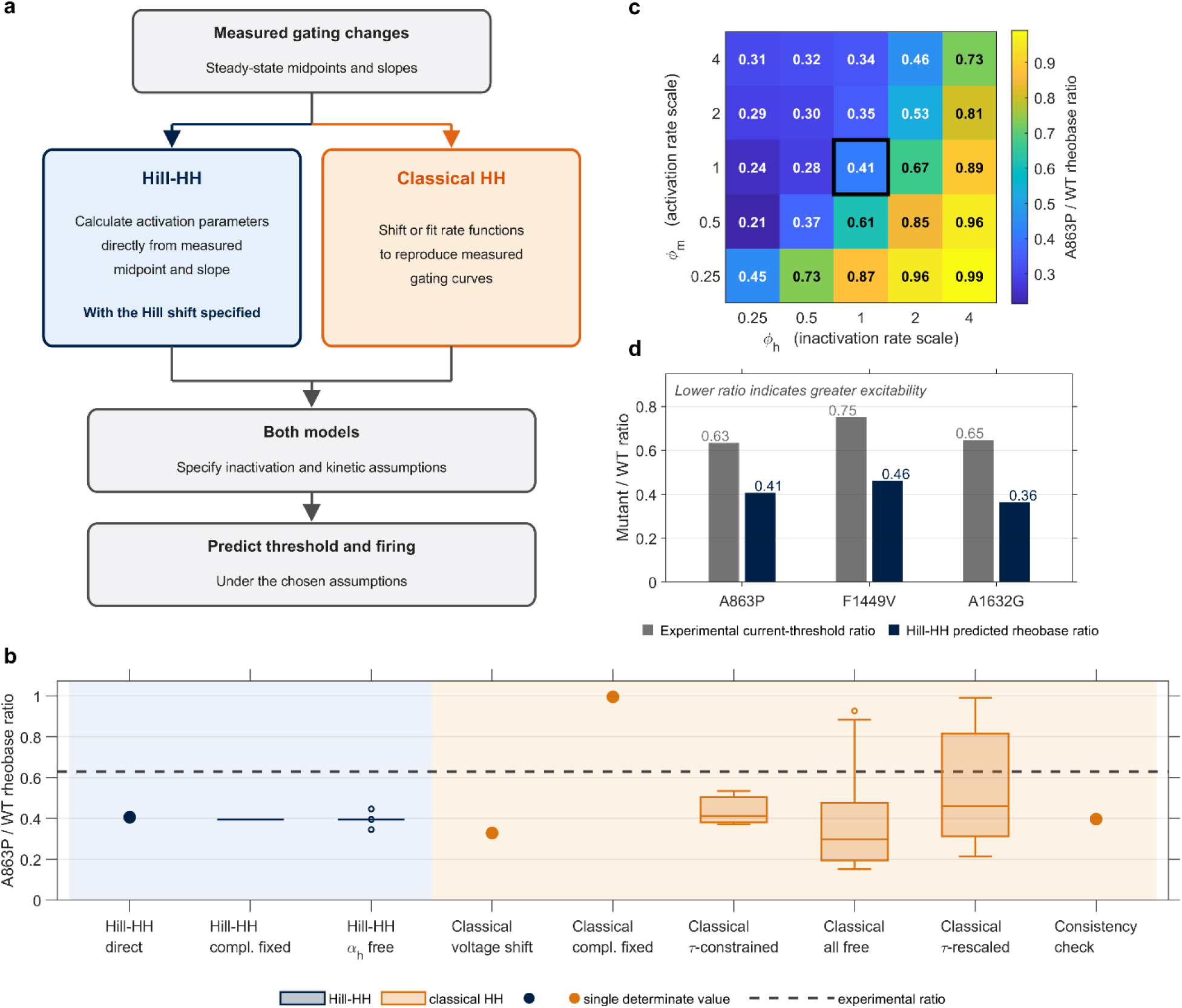
Mutation mapping and sensitivity to kinetic assumptions. **a**, Schematic of the mapping from reported steady-state gating descriptors to model implementations. The Hill-based mapping retains specified complementary rates and an assumed inactivation construction; steady-state descriptors alone do not determine gating kinetics in either formulation. **b**, A863P-to-wild-type (WT) rheobase ratios across Hill-based and classical Hodgkin-Huxley (HH) implementations. The direct Hill implementation maps the selected gating descriptors to a single parameter set. Hill ensembles either retain the complementary rates or additionally allow the inactivation rate αh to vary. Classical HH implementations comprise fits with all rate parameters free or complementary rates fixed, voltage shifts of the reference rate functions, fits constrained by assumed time constants, and a kinetic-rescaling grid. The consistency check fits classical HH rates to Hill-derived inactivation and therefore tests agreement between implementations rather than independent predictive performance. In each box the central line is the median, the box spans the 25th to 75th percentiles, the whiskers extend to the most extreme values within 1.5 times the interquartile range of the box edges, and values beyond that are plotted individually as outlier markers. Implementations yielding a single estimate are shown as points without a box. The dashed line indicates the experimental mutant-to-WT current-threshold ratio (approximately 0.63). Implementation definitions and numerical summaries are provided in Supplementary Table 7. **c**, Classical HH rheobase ratios after multiplying both forward and backward activation rates by ∅*_m_* and both inactivation rates by ∅_ℎ_. This scaling preserves the steady-state gating curves while dividing the corresponding time constants by the scaling factor. The outlined cell denotes the unscaled reference (∅*_m_* = ∅_ℎ_ = 1). **d**, Experimental mutant-to-WT current-threshold ratios compared with direct Hill-model rheobase ratios for A863P, F1449V, and A1632G (Supplementary Table 12). Both quantities describe relative threshold changes within their respective experimental or model settings; lower ratios indicate a greater threshold reduction. The Hill-based predictions reproduce the direction of the experimental changes but overestimate their magnitude. Representative voltage responses at an applied current of 40 µA/cm² are shown in Supplementary Figures 3 and 5.

The same conclusion was evident in the representative voltage traces (Supplementary Figure 3c,g) and firing-frequency curves (Supplementary Figure 3d,h). At comparable applied currents, the A863P-inspired condition entered repetitive firing at lower current and reached substantially higher firing rates than the WT condition in both implementations of this consistency comparison, while spike amplitude remained broadly preserved. This agrees qualitatively with the experimental observation that A863P-expressing DRG neurons show reduced current threshold and increased firing frequency without a significant change in spike amplitude^34^. The simulated rheobase reduction was larger than that reported experimentally. The benchmark does not include Nav1.8 and other DRG-specific conductances; their contribution to this discrepancy was not tested.

Importantly, the case study does not indicate that the Hill-based formulation predicts the A863P phenotype more accurately than the original HH consistency comparator. Instead, it shows that the reformulated model preserves the classical framework’s ability to represent a biologically motivated sodium-channel perturbation, while expressing the required gating changes through more transparent and functionally interpretable parameters. This directly addresses the practical motivation for the reformulation: making experimentally reported gating shifts easier to encode, inspect, and tune within a conductance-based model.

### 3.6 A863P predictions depend on kinetic assumptions

Direct Hill mapping gave WT and A863P rheobases of 17.69 and 7.19 µA/cm², respectively (ratio 0.406), compared with the experimental current-threshold ratio of 0.633. At 40 µA/cm², WT fired once and A863P fired 16 spikes during the 100-ms step (Figure 5b; Supplementary Table 7). The activation-only and inactivation-only ratios were 0.459 and 0.919 (Supplementary Table 11).

With complementary rates fixed, all 200 random-start Hill fits were accepted and returned the same rheobase ratio of 0.395, with one WT spike and 16 mutant spikes at 40 µA/cm². When the amplitude and voltage scale of the inactivation forward rate were released, all 200 fits were again accepted; ratios ranged from 0.346 to 0.446 (median 0.395). A863P fired 15-16 spikes in every fit, while WT fired once in 196 of 200 fits (Figure 5b; Supplementary Figure 6b; Supplementary Table 7). Peak inactivation time constants ranged from approximately 1.2 to 32 ms, compared with roughly 30-145 ms in the classical ensembles across the accepted fits (Supplementary Figure 4 shows 80 fits per panel on a logarithmic axis for clarity).

These Hill ensembles used the Boltzmann activation target but retained the inactivation curve chosen in the direct Hill implementation. Their repeatability is therefore conditional on the selected inactivation construction and complementary rates. Both formulations require kinetic assumptions when only steady-state measurements are available (Figure 5a; Supplementary Table 7).

When all classical sodium-rate parameters were fitted to the published Boltzmann targets, all 200 starts met the acceptance criterion. Every accepted fit predicted a lower mutant rheobase, but ratios ranged from 0.15 to 0.93 despite agreement with the same steady-state targets (Figure 5b; Supplementary Figure 6a; Supplementary Table 7). The ensemble median ratio was 0.30. Thus, the direction of the threshold change was retained, whereas its magnitude was not identified by the steady-state fits.

Fixing the complementary classical rates produced a different outcome. Only 104 of 200 starts were accepted, and all gave WT and mutant rheobases of 47.2 and 47.0 µA/cm², respectively (ratio 0.996). This was a negligible reduction, rather than the substantial experimental effect. Both conditions were silent at 40 µA/cm². A voltage-shift-only implementation instead gave a ratio of 0.33, but retained classical slopes rather than matching the full Boltzmann targets; WT and mutant each fired one spike at 40 µA/cm² (Figure 5b; Supplementary Table 7).

Constraining the time constants near voltage-shifted classical references gave a ratio of 0.413 at the reference penalty weight of 0.05, with rheobases of 39.16 and 16.16 µA/cm². Maximum steady-state error was at most 0.012, and time constants remained within 0.72-1.30 times their reference values. Changing the penalty weight from 0.1 to 0.02 moved the ratio from approximately 0.37 to 0.54 (Figure 5b; Supplementary Table 7). The inferred threshold change therefore depended on a kinetic constraint not supplied by the experimental steady-state measurements.

Rescaling the activation and inactivation rates around the reference fit isolated this dependence. The 25 combinations retained identical steady-state curves but produced rheobase ratios from 0.21 to 0.99; the unscaled reference gave 0.413 (Figure 5c). All ratios remained below one, and the range spanned the experimental ratio of 0.633. This span demonstrates sensitivity to unconstrained kinetics; it does not establish an exact match to the experimental ratio at a sampled grid point. Neither the rescaling grid nor the all-free and time-constant-constrained classical fits reproduced sustained repetitive mutant firing at 40 µA/cm² (Supplementary Table 7).

The classical consistency comparator described above gave a ratio of 0.397, close to the direct Hill value (Figure 5b; Supplementary Figure 3). Because its inactivation target was a Hill-model output, this agreement is a consistency check rather than an independent prediction from the published inactivation curve.

### 3.7 Three mutation benchmarks reproduce qualitative changes but overestimate threshold reductions

We applied the direct Hill mapping to F1449V^51^ and A1632G^52^ using study-specific WT controls (Supplementary Table 8). F1449V lowered rheobase from 17.31 to 7.97 µA/cm², giving a ratio of 0.460 compared with the experimental ratio of 0.750. A1632G lowered rheobase from 13.78 to 5.00 µA/cm², giving a ratio of 0.363 compared with 0.646 experimentally (Figure 5d; Supplementary Tables 11 and 12). Activation-only ratios were 0.505 and 0.424, whereas inactivation-only ratios were 0.913 and 0.900, respectively (Supplementary Table 11).

Both mutants fired repetitively at 40 µA/cm², whereas their respective WT models fired once; frequency-current curves also showed earlier firing and higher discharge rates in the mutant conditions (Supplementary Figure 5). First-spike voltage thresholds were nearly unchanged. Width changed from 1.6 to 1.7 ms for F1449V and remained 1.3 ms for A1632G, while amplitudes increased by approximately 6% and 10%, respectively. AHP was more pronounced in both mutants (Supplementary Table 13). These deterministic waveform comparisons do not establish statistical equivalence to experimental recordings.

Across A863P, F1449V, and A1632G, direct Hill ratios ranged from 0.363 to 0.460, compared with experimental ratios of 0.633-0.750 (Figure 5d). The mapping reproduced lower threshold and enhanced repetitive firing, but consistently overestimated the threshold reduction. These Hill-only extensions broaden the qualitative benchmark; they do not establish superiority over classical HH across mutations. Together with the A863P sensitivity analyses, they separate repeatable mapping under stated assumptions from quantitatively accurate prediction of neuronal phenotype.

## Discussion

The present study introduces a Hill-based reformulation of selected Hodgkin-Huxley (HH) kinetic terms and shows that this modification preserves the core dynamical behavior of the classical HH model while reorganizing its parameter space into an explicit set of Hill control axes^2^. The central result is not simply that Hill functions can fit the historical HH-derived relationships, but that this choice produces a useful hybrid conductance-based model in which steady-state gating remains close to the original formulation, canonical excitability is retained, experimentally motivated gating shifts can be encoded directly, and the mapping from kinetic parameters to spike-waveform phenotype can be inspected along these coordinates ^1,2,12,31,36,37,37^.

Several strategies have sought to overcome limitations of the original HH rate expressions. Reduced and phenomenological models improve tractability or reproduce broad firing repertoires, but they weaken the direct link to ionic gating ^7–9^. Boltzmann parameterizations retain that link and are widely used in voltage-clamp fitting and multicompartment modeling ^15–17^. The present study asks whether a three-parameter Hill family can provide a compact alternative description of the same HH-derived gating targets ^1,7,9–12,27,53,54^.

Among the compact sigmoidal families tested here - Hill, Boltzmann, hyperbolic tangent, error function, and arctangent - the Hill function yielded the lowest SSE across the three original HH-derived targets, acceptable residual behavior by the Anderson-Darling (AD) test, and overall competitive AIC values^55–57^ (Methods). This is notable because these targets span distinct voltage-dependence profiles, from the relatively gradual potassium-activation relationship to the much steeper sodium-activation and sodium-inactivation transitions. The Hill form therefore provides a compact and flexible description of the selected gating nonlinearities while retaining interpretable parameters corresponding to horizontal shift, Hill midpoint, and steepness. For inactivation, the Hill midpoint describes the rate function rather than the derived steady-state curve. Whether this parameterization extends meaningfully to the fitting of experimental recordings from diverse neuron types remains to be demonstrated, but the present benchmark establishes the empirical foundation for that hypothesis ^20,22,25,55–57^.

A second important result is that the reformulation is conservative where it needs to be and selective where it can be informative. The comparison of derived gating quantities shows that the modified model preserves the steady-state activation and inactivation structure of the original HH formulation to a large extent, particularly for *m*_∞_(*V*), while introducing more substantial changes in the time constants, especially *τ*_ℎ_(*V*) and *τ_n_*(*V*). This distinction is functionally informative: the Hill-based reformulation leaves intact the voltage ranges over which the major gating processes are recruited, but reshapes the timescales over which those processes unfold. That asymmetry offers a plausible explanation for why the modified model retains canonical HH-like spike waveforms and *f*–*I* structure while showing a modest shift in effective excitability. The current-remapping analysis makes this concrete: the principal difference between the original and modified models is a smooth, approximately affine shift in the applied current required to recover matched responses, with R² = 0.99 over the explored range. The Hill-based substitution retains the firing patterns examined here; it preserves the canonical dynamical phenotype while shifting the effective operating point along the input axis. This is exactly the desired behavior for a reparameterization intended to improve interpretability without sacrificing the conductance-based logic of the original model ^1,2,23,26,40^.

Beyond preserving classical HH dynamics, the reformulated model also changes how excitability is organized in parameter space. The two-dimensional regime maps show that the Hill parameters partition the *I_app_* − *θ* plane into broad, contiguous domains corresponding to silence, single spiking, tonic driven firing, spontaneous activity, and depolarization block. This offers a practical way to inspect parameter effects; in the classical empirical HH parameterization, tuning the kinetics typically involves manipulating several constants whose dynamical roles are indirect and coupled. In the Hill-based formulation, parameters grouped by functional meaning - shift, Hill midpoint, and steepness - organize the observed transitions between regimes. Activation-related parameters strongly influence the onset of spontaneous activity, whereas inactivation-related parameters more prominently shape the persistence of driven firing and the transition into depolarization block. This is worth comparing explicitly with the Boltzmann-based literature: while Boltzmann parameters likewise have half-activation and slope interpretations, the three-parameter Hill formulation adds a horizontal-shift parameter alongside midpoint and steepness. Whether this additional degree of freedom translates into practical advantages for fitting or navigation in specific cell types is an open question that the present parameter-space maps motivate ^30–33,39^.

The spike-feature heat maps sharpen this point further. Across the biologically admissible parameter ranges explored here, the modified model preserves control over amplitude, threshold, width, and AHP while varying nine Hill coordinates. In several panels, especially those associated with sodium activation and potassium activation, parameter-to-feature relationships appear more orderly and easier to interpret than in the original HH parameterization, where comparable effects are dispersed across a larger set of empirical constants. These results support the practical value of the Hill coordinate system for inspecting how specific gating changes reshape waveform properties, while leaving formal range-normalized comparisons of landscape smoothness and coverage for future work.

These features are especially relevant in the broader context of conductance-based modeling. Classical HH-type models remain central because they offer a biophysically and computationally tractable description of excitability, but their historical empirical rate laws are not always the most convenient basis for adapting the framework across cell types, preparations, or modeling goals. More detailed channel-state or Markov models can offer richer biophysical realism at the cost of substantially greater dimensionality and reduced practical identifiability ^58–60^. Boltzmann-based parameterizations occupy a useful middle ground, and the Hill formulation offers an alternative coordinate system; the present analyses do not establish superior prediction over Boltzmann-based models. The Hill-based reformulation explored here occupies a complementary position: it retains the essential conductance-based architecture of HH, preserves much of its canonical dynamical behavior, and yet provides a parameterization with nine tunable Hill coordinates that is explicitly grounded in a biologically motivated functional family, with the remaining classical rates retained ^1,2,27,28,33,61^.

The Nav1.7-A863P benchmark provides a biologically motivated illustration of this practical value. Both the original HH consistency comparator and the Hill-based model reproduced the expected qualitative consequence of the A863P gating perturbation: reduced rheobase and enhanced repetitive firing. The two implementations behaved similarly at the output level for this shared inactivation target, which argues against presenting the Hill-based model as more accurate in this case. Instead, the useful distinction is representational. The Hill formulation encoded the reported activation and inactivation shifts through explicit gating parameters, whereas the original HH consistency comparator required coordinated adjustment of empirical rate constants to reproduce the activation targets and the Hill-derived inactivation curve. This is the intended use of the reformulation: preserving HH-like dynamics while making experimentally reported channel perturbations easier to encode and interpret^73^.

The broader A863P comparisons show why that distinction matters. Classical fits to the published Boltzmann steady-state targets gave rheobase ratios of 0.15-0.93 across accepted all-free fits, whereas fixing the complementary rates gave almost no threshold change (Figure 5b; Supplementary Figure 6a; Supplementary Table 7). Rescaling time constants while preserving the same steady-state curves produced ratios of 0.21-0.99 (Figure 5c). The experimental ratio lies within this range, but the range alone does not identify the appropriate kinetics. Agreement with a steady-state curve is therefore insufficient to determine either the magnitude of the threshold change or the firing response.

The Hill ensembles were more repeatable within the tested construction, but their inactivation target was the curve selected in the direct Hill implementation rather than the published Boltzmann curve. Retaining classical complementary rates, or releasing selected parameters of the inactivation forward rate, also restricts the possible kinetics (Supplementary Figures 4 and 6b). These choices are assumptions in both formulations. The narrow Hill ensemble therefore supports reproducibility for the specified mapping, rather than resolving the kinetic information missing from the measurements.

F1449V and A1632G extended the direct Hill benchmark beyond A863P. Across the three mutations, predicted mutant/WT rheobase ratios ranged from 0.363 to 0.460, compared with experimental current-threshold ratios of 0.633-0.750 (Figure 5d; Supplementary Tables 11 and 12). The model reproduced lower threshold and enhanced repetitive firing, but consistently overestimated the threshold reduction. First-spike features provided an additional comparison for F1449V and A1632G (Supplementary Table 13), although deterministic model differences cannot establish statistical equivalence to recordings. These retrospective Hill-only extensions support qualitative transfer of the mapping across mutations, not a general quantitative advantage over classical HH.

Several limitations should be made explicit. First, the present model is deliberately hybrid: it recasts selected sigmoidal kinetic terms while retaining the remaining classical components. The conclusions therefore support this specific hybrid reformulation rather than a universal argument that Hill functions should replace all HH-type rate laws. Second, both the primary fitting analysis and most simulations remain anchored to the original squid-axon dataset at 6.3°C. The Nav1.7-A863P, F1449V, and A1632G case studies add biologically motivated perturbation benchmarks, but the model is not a full DRG-neuron reconstruction because it does not include Nav1.8 and other sensory-neuron conductances. Agreement with the experimental phenotypes should therefore be interpreted qualitatively: the direct Hill mappings reproduce the direction of the threshold and firing-frequency changes, but overestimate the magnitude of the threshold. Classical A863P outcomes vary with the implementation (Figure 5; Supplementary Tables 7-12). The contribution of the omitted conductances to these discrepancies was not tested ^34^. Third, slope factors for F1449V and A1632G were recovered from published curves rather than directly reported, adding uncertainty to the input descriptors (Supplementary Table 8). The Hill coefficient is used here as a phenomenological steepness parameter and should not be interpreted as direct evidence of molecular cooperativity in voltage-gated channels. Finally, formal bifurcation and continuation analyses, temperature-dependent scaling, and fitting to voltage-clamp datasets from identified neuron types remain important next steps^62–64^.

Despite these limitations, the present results support a clear conclusion. The Hill-based reformulation introduced here preserves the essential excitability structure of the classical HH model while replacing part of its historical empirical parameterization with an explicit set of Hill control variables. The resulting model is not a universal replacement for the classical HH formalism, but a complementary parameterization for fitting, perturbing, and interpreting selected gating relationships. Its contribution is a reproducible mapping under stated assumptions, rather than demonstrated superiority in quantitative prediction. For conductance-based modeling, this combination of fidelity, compactness, and interpretability is valuable, and suggests that the Hill-based HH reformulation may serve as a useful complement to existing parameterization strategies for exploring cell-specific ion-channel dynamics ^1,2,20,22,25,36,45^.

## Methods

### Classical Hodgkin-Huxley Model

We used the classical HH model as the reference conductance-based description of action-potential generation in an excitable membrane. In this formulation, the membrane is represented as a capacitance in parallel with voltage-dependent sodium and potassium conductances and a passive leak current. The corresponding current-balance equation is ^1–6,65,66^

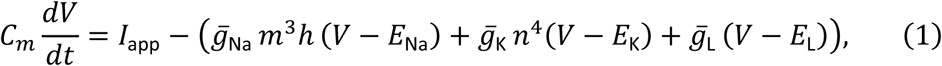

where *V* is membrane potential, *I*_app_ is the externally applied current, *C_m_* is membrane capacitance per unit area, *g̅*_Na_, *g̅*_K_, *g̅*_L_ are the maximal sodium, potassium, and leak conductances, and *E*_Na_, *E*_K_, *E*_L_ are the corresponding reversal potentials. Sodium activation, sodium inactivation, and potassium activation are governed by the gating variables *m*, ℎ, and *n* respectively. Each gating variable evolves according to first-order voltage-dependent kinetics of the form

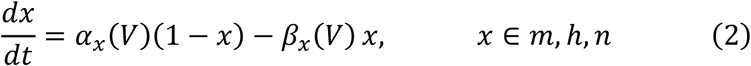

which can equivalently be written in relaxation form as

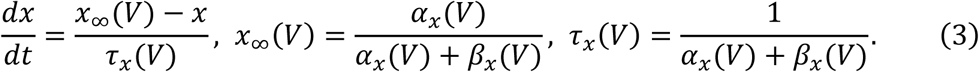

The classical HH voltage-dependent rate functions were used in their original form. Simulations were performed under the standard squid-axon conditions used for benchmarking in this study. The membrane potential was initialized at the resting state, and the gating variables were initialized at their corresponding starting values from the classical HH formulation. Temperature-dependent scaling factors were not included in the present work, so all comparisons between the original and modified models were performed under the same unscaled baseline conditions^1,2,6,67^.

Under voltage-clamp steps to fixed potentials, these equations predict exponential convergence of each gating variable to its steady-state value with time constant *τ_x_*(*V*), providing a direct link between measured current transients and inferred gating curves ^1,2^. In the original HH model, the voltage-dependent forms of *α_x_*(*V*) and *β_x_*(*V*) were chosen empirically to fit squid giant axon data and, together with the exponents in the sodium and potassium conductances, define the classical parameterization of membrane excitability. Specifically, the HH rate functions were fitted using exponential-rational forms, with voltage expressed in mV and time in ms. Figure 6 reproduces the canonical voltage dependence of these rate functions as reported by Hodgkin and Huxley in 1952. Related parameterizations have since been widely adopted and modified to reproduce neuronal behaviors across species and cell types ^13,68,69^. The original HH rate parameter values are summarized in Supplementary Table 1 ^1–6,70,71^.

**Figure 6.**
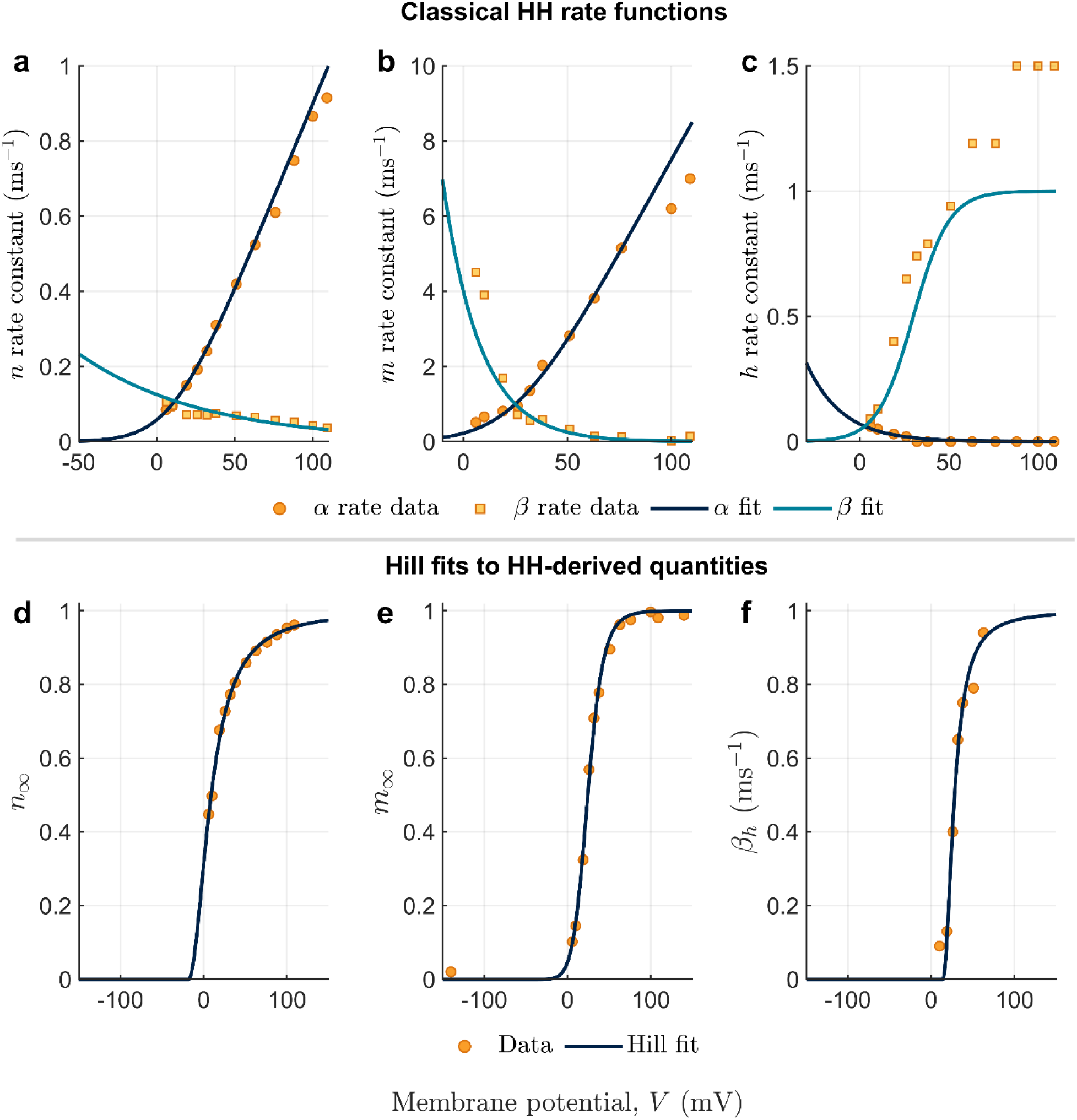
Classical Hodgkin-Huxley voltage dependence and Hill-type fits to the original Hodgkin-Huxley dataset. **(a-c)** Canonical Hodgkin-Huxley gating kinetics plotted as functions of membrane potential using the original experimental measurements reported by Hodgkin and Huxley. Panels **a-c** correspond to the *n*, *m*, and ℎ gating variables, respectively. Circular markers denote experimentally derived *α* rate constants, square markers denote experimentally derived *β* rate constants, navy curves denote the classical Hodgkin-Huxley *α* functions, and blue curves denote the classical Hodgkin-Huxley *β* functions. Here, *α* and *β* represent the voltage-dependent forward and backward gating rate functions in the Hodgkin-Huxley formalism. **(d-f)** Hill-type fits obtained from quantities derived from that same original dataset and used in the modified model, namely *n*_∞_(*V*), *m*_∞_(*V*), and *β*_ℎ_(*V*). Orange markers indicate the original data and navy curves the corresponding Hill fits. In all three cases, the Hill functions closely reproduce the empirical voltage dependence, capturing both the curvature and asymptotic saturation of the original measurements. Despite differences in steepness across variables, the fitted profiles preserve the distinct kinetic signatures of the underlying gating processes, including the more gradual voltage dependence of potassium activation and the sharper transitions associated with sodium activation and inactivation. Together, these results show that selected components of the original Hodgkin-Huxley formalism can be recast using a compact Hill-based parameterization while remaining anchored to the original experimental data.

However, the original HH formulation defines membrane voltage as a displacement from the resting membrane potential using a sign convention opposite to that commonly adopted in modern electrophysiology. Under the original convention, depolarization corresponds to positive voltage displacements from rest, whereas modern membrane-potential representations typically assign more positive values to membrane depolarization relative to an absolute resting potential. To maintain consistency with contemporary voltage-polarity conventions and facilitate interpretation of the fitted voltage-dependent relationships, all HH voltage-dependent equations were reformulated using the negated variable (−V). The same transformation was applied to the experimental HH-derived data used for fitting in the next subsection^2,6,65^.

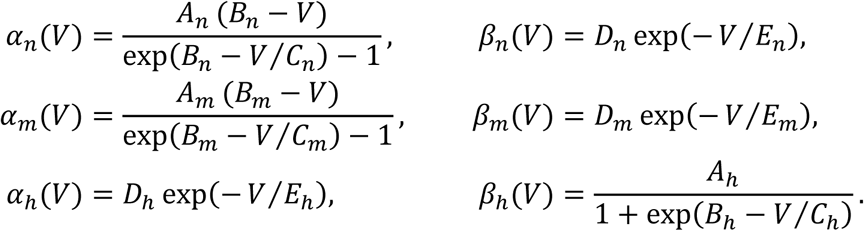

Although the HH framework generalizes broadly, the specific forms used for *α_x_*(*V*) and *β_x_*(*V*) can become restrictive when other preparations exhibit steeper, more saturating, or otherwise preparation-specific voltage dependences. Motivated by this, and by the central role of *m*, ℎ, and *n* in shaping excitability, we replace selected classical expressions with Hill-type sigmoidal parameterizations fitted directly to HH-derived voltage-dependent data. The following subsections describe the adopted fitting forms, the preprocessing applied to the historical dataset, and the criteria used to evaluate the substitutions ^1,23,26,27,29,61^.

Unless otherwise stated, all simulations use the classical HH baseline at 6.3 °C, as summarized in Supplementary Table 2. This temperature matches the conditions of the original Hodgkin-Huxley voltage-clamp experiments and therefore provides an internally consistent reference for evaluating the Hill-based reformulation. Extension of the present framework to other temperatures or preparations would require corresponding voltage-clamp data acquired under those conditions ^2,6,67^.

The membrane potential is initialized to −65 mV and the gating variables are initialized as ℎ = 0.81, *n* = 0.32 and *m* = 0.04. Currents and conductances are expressed per unit area, with capacitance given in *μ*F/cm^2^. Temperature-dependent scaling effects (e.g., *Q*_10_ adjustments of rate constants) are not included. Supplementary Table 2 therefore serves as the benchmarking baseline against which we evaluate the impact of replacing selected classical rate expressions with the proposed Hill-type parameterizations.

### Model Modifications Using Sigmoidal Fits

Among the six classical HH kinetic relationships, only *m*_∞_(*V*), *n*_∞_(*V*), and *β*_ℎ_(*V*) exhibit monotonic, saturating sigmoidal profiles over the physiological voltage range. We therefore selected these three quantities for sigmoidal substitution and retained the remaining rate functions in their classical HH form. The remaining three - *β_n_*(*V*), *β_m_*(*V*), and *α*_ℎ_(*V*) - do not: *β_n_*(*V*) and *β_m_*(*V*), are monotonically decreasing exponentials with no saturation plateau, and *α*_ℎ_(*V*) is likewise a pure exponential without sigmoidal structure. These functional forms are therefore not natural candidates for a Hill-type substitution, which is designed specifically for steep saturating input-output relationships. To avoid introducing unnecessary complexity, and because steady-state quantities are more directly interpretable and experimentally accessible than the raw forward and backward rates individually, we further expressed the sodium and potassium activation as steady-state curves *m*_∞_(*V*) and *n*_∞_(*V*) rather than fitting *α_m_*(*V*) and *α_n_*(*V*) directly. The sodium inactivation rate *β*_ℎ_(*V*) was fit directly rather than through ℎ_∞_(*V*) because *β*_ℎ_(*V*) itself has the requisite sigmoidal shape and direct fitting avoids an unnecessary algebraic transformation. Together, these choices define the minimal Hill-based substitution that replaces every HH component with a sigmoidal voltage dependence.

To confirm that this selection is grounded in the original experimental substrate, we returned directly to the Hodgkin-Huxley dataset and examined whether the target gating relationships could be recast using a compact Hill-type formalism. Figure 6 is built from the same historical measurements reported by Hodgkin and Huxley themselves, rather than from synthetic data or from refitting a different preparation. For consistency with contemporary electrophysiological voltage conventions, the original Hodgkin-Huxley voltage values were transformed using (−V) prior to analysis and fitting; however, the underlying experimental measurements remained unchanged. Panels a-c reproduce the canonical experimental data underlying the original rate functions and illustrate the distinct kinetic profiles that motivate the reformulation: potassium activation displays a comparatively gradual voltage dependence, whereas sodium activation and sodium inactivation exhibit sharper and more strongly nonlinear transitions. These differences confirm that the selected components of the classical HH formalism can be represented by steep but saturating sigmoidal functions without sacrificing their characteristic kinetic signatures ^1–5,23,26^.

Panels d-f show the Hill-type fits to *n*_∞_(*V*), *m*_∞_(*V*), and *β*_ℎ_(*V*), performed on quantities derived from that same historical dataset. The resulting fits closely reproduce the original HH data across the voltage range examined, capturing both the curvature and the asymptotic behavior of the empirical measurements. The potassium activation curve *n*_∞_(*V*) is well captured with a relatively low Hill coefficient, consistent with its gradual voltage dependence, while *m*_∞_(*V*) and *β*_ℎ_(*V*) require substantially higher coefficients, reflecting the sharp voltage sensitivity associated with sodium activation and inactivation. This reformulation introduces three parameters per fitted variable - horizontal shift a, half-(in)activation voltage *V*_ℎ*alf*_, and Hill coefficient l - each with direct biological interpretation that can be tuned independently, replacing the corresponding empirical rate laws with a more transparent functional description ^1–5,20,22,25^.

The fitted data points were taken from the original Hodgkin–Huxley study using −*V* to match the sign convention adopted throughout this work, and are summarized in Supplementary Table 3. Parameter estimation was performed with MATLAB’s lsqcurvefit (Optimization Toolbox) by minimizing the sum of squared errors (SSE) between each candidate sigmoidal function and the historical HH-derived data ^72^.

For the *β*_ℎ_(*V*) fit specifically, two auxiliary data points were introduced at the low- and high-voltage extremes to constrain the asymptotic behavior of the sigmoid, because the original historical dataset is sparse near saturation. These auxiliary points were used only to stabilize the fit in the head and tail regions of the curve. In addition, repeated values greater than 1 were removed from the historical *β*_ℎ_(*V*) table before fitting. These preprocessing steps were applied consistently across all candidate sigmoidal families and should be interpreted as pragmatic adjustments to stabilize the fit to the historical HH-derived dataset rather than as a redefinition of the underlying HH formalism. To ensure that these preprocessing choices did not drive the model-selection result, the fits were repeated with the original dataset retained and the auxiliary asymptotic points omitted; the relative advantage of the Hill fit was preserved^55,73^.We compared five compact sigmoidal families - Boltzmann, arctangent, hyperbolic tangent (tanh), error function (erf), and Hill function - using the same datasets and optimization routine. Model performance was evaluated using the Anderson-Darling (AD) test on residuals, the sum of squared errors (SSE), and the Akaike information criterion (AIC) ^55–57^. Supplementary Table 4 summarizes these comparisons. The AD test was used to assess whether the residuals were consistent with the assumed distributional behavior, namely normality. For each fit, residuals were computed as the difference between the HH-derived data and the fitted function at the same voltages, standardized by subtracting their mean and dividing by their standard deviation, and tested with MATLAB’s adtest. A p-value below 0.05 was taken as evidence of departure from normality, whereas a p-value above 0.05 indicated only a failure to reject normality and was not treated as confirmation of fit adequacy. SSE quantified absolute fitting error and AIC provided a complexity-penalized measure of goodness of fit, computed in its least-squares form as *AIC* = *Nln*(*SSE*/*N*) + 2*k*. Here *N* is the number of fitted data points and k is the number of free parameters: *k* = 2 for the Boltzmann, tanh, and erf functions and *k* = 3 for the arctangent and Hill functions. AIC values were compared among fits to the same target data, with lower values indicating greater relative support. These metrics were used jointly to select the sigmoidal family adopted in the modified model, balancing numerical accuracy, statistical adequacy, and parameter parsimony.

Because visual agreement alone does not distinguish among compact sigmoidal families, the same HH-derived datasets for *n*_∞_(*V*), *m*_∞_(*V*), and *β*_ℎ_(*V*) were fit with the Hill, Boltzmann, tanh, erf, and arctan formulations. This side-by-side comparison, summarized in Figure 7 and Supplementary Table 4, was used to justify the final choice of fitting family. All five candidate functions captured the broad sigmoidal shape of the target relationships, but their quantitative agreement with the HH-derived data differed. The Hill family was therefore selected for the modified formulation on the basis of its overall performance across the three fitted targets ^55–57^.

**Figure 7.**
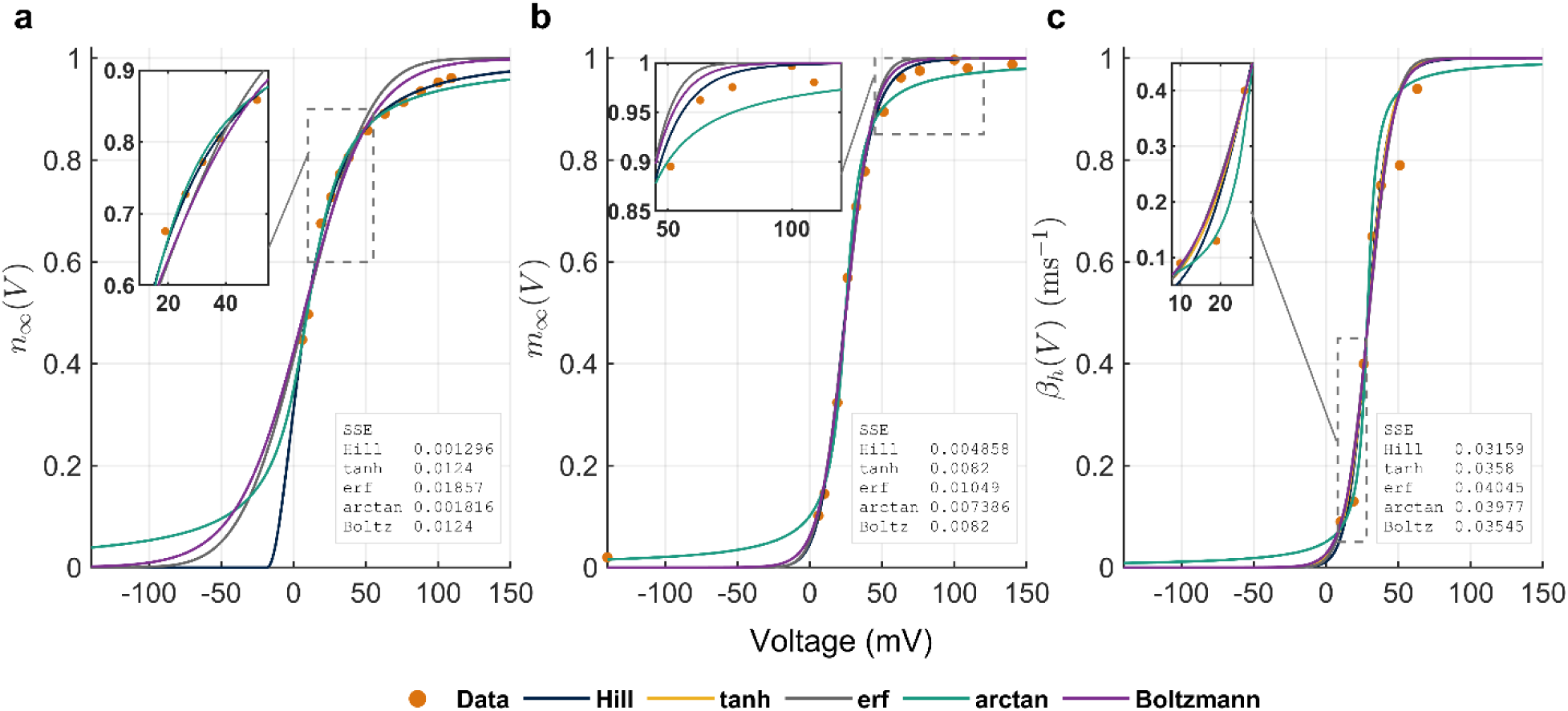
Comparison of compact sigmoidal fits to Hodgkin-Huxley-derived gating data. Fits of Hill, Boltzmann, tanh, erf, and arctan functions to the original Hodgkin-Huxley-derived relationships **(a)** *n*_∞_(*V*), **(b)** *m*_∞_(*V*), and **(c)** *β*_ℎ_(*V*). Orange markers indicate the data, and colored curves the fitted functions. Insets provide zoomed view of selected regions highlighting local differences between the fitted functions. The boxes within each panel report the corresponding sum of squared error (SSE) of each fit. Across all three targets, the Hill function yields the lowest fitting error, indicating the best overall agreement with the historical Hodgkin-Huxley data and justifying its selection for the modified model.

The choice of the Hill family was motivated not only by fitting accuracy but also by interpretability. In the Hill parameterization, the fitted coefficients map naturally onto horizontal shift, Hill midpoint voltage, and steepness, making the resulting parameter space easier to relate to channel-gating behavior than a purely generic sigmoid fit. Accordingly, the Hill function was adopted as the compact reparameterization used in all subsequent analyses of the modified model ^19,20,22,25^.

As a result, we employ the Hill function as a sigmoidal fitting function, ranging in [0,1] and taking the following form ^20,22^:

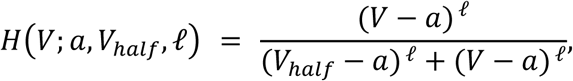

where *a* is a horizontal shift, *V*_ℎ*alf*_ sets the midpoint of the fitted Hill relationship, and l > 0 is the Hill coefficient that is responsible for the slope or steepness. To robustly handle negative values when l is non-integer during the optimization process, we evaluate powers via a sign-safe formulation:

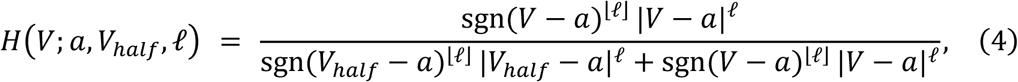

with sgn(⋅) the sign function and ⌊l⌋ the integer part of l. Following the optimization, we set *H*(*V*) = 0 for *V* < *a* to enforce the desired S shape.

For comparison, the following sigmoidal forms were used for the remaining functions, where a, *V*_0_, and *k* represent the free parameters during the optimization fitting process ^7,9–11^:

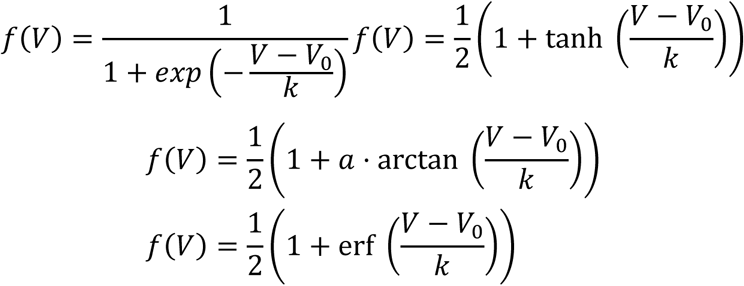

We fit equation (4) to the experimental data points summarized in Supplementary Table 3:

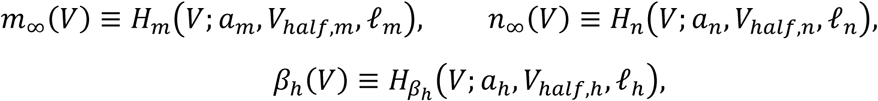

The fitted Hill expressions for *n*_∞_, *m*_∞_ and *β*_ℎ_ obtained by minimizing the sum of squared errors, are:

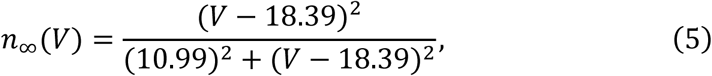

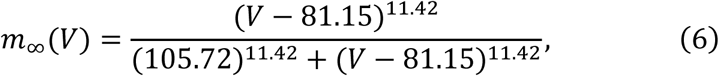

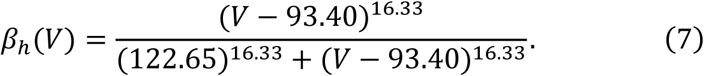

The remaining rates are kept as in the classical HH paper:

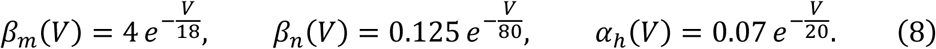

Given the steady-state identity *x*_∞_(*V*) = *α_x_*(*V*)/(*α_x_*(*V*) + *β_x_*(*V*)), we compute the *opening* rates from the fitted *m*_∞_ and *n*_∞_ given by equations (6) and (5), respectively

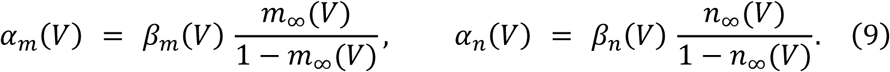

For the inactivation gate ℎ, we directly use the fitted *β*_ℎ_(*V*) given by equation (7) together with the original *α*_ℎ_(*V*). The resulting parameterization enters the same dynamical system defined above, with the fitted activation relationships used to derive the corresponding opening rates and with βh replaced by the fitted Hill expression.

The parameter ranges used in all analyses were chosen to keep the model within physiologically meaningful and biologically realistic firing regimes. Here, physiologically meaningful values were defined by Hill coefficients greater than 2 and shift parameters lower than their corresponding half-(in)activation voltages. Biologically realistic firing was defined as non-damping spike trains containing at least three spikes with peak voltages above 0 mV. These ranges were used for the one-parameter heat-map sweeps and were expanded by 15% on both sides, when applicable, to generate the two-dimensional excitability maps. The final ranges explored for the modified and classical HH models are summarized in Supplementary Tables 5 and 6 ^30–33,39^.

For the baseline Hill-model parameter sweeps, parameters appearing only in retained classical HH rate functions were held fixed and excluded from the tunable parameter set. This restriction does not apply to the A863P kinetic-sensitivity ensembles described below, in which selected complementary-rate parameters were explicitly released.

### Simulation Protocols, Spike Detection, and Electrophysiological Feature Analysis

All simulations were implemented in MATLAB R2024b using the numerical solver ode113. Unless otherwise stated, the models were simulated for 200 ms with temporal resolution of 0.01 ms and constant applied current injected between 50 and 150 ms. The applied current *I_app_* was expressed in (*μ*A/cm^2^), consistent with the units used in the original HH formulation. Spike detection was performed using an amplitude-based algorithm, where only spikes exceeding 40% of the maximum membrane potential attained within the trace were identified.

For firing frequency calculations of Figure 2b, applied current was varied from 0 to 200 *μ*A/cm^2^ in increments of 5 and for each *I_app_* value the number of detected spikes during the simulation was divided by the time interval to obtain the firing rate in Hz.

For the excitability maps shown in Figure 3, the corresponding parameter ranges were uniformly divided into 15 values, where each simulated trace was classified into one of the five activity regimes based on its spike pattern. A trace was classified as silent if no spikes were detected during the simulation. It was classified as single spike if exactly one spike occurred during the stimulation interval (50-150 ms), and no more than one spike occurred when no current was applied. Tonic firing was defined as the occurrence of two or more spikes during stimulation interval, and no more than one spike outside this interval. Spontaneous firing was assigned when two or more spikes occurred without stimulating current, in addition to at least one spike within stimulation interval. Depolarization block was identified when spiking activity was initiated during stimulation but failed to develop into sustained tonic firing, instead the membrane potential remained persistently depolarized above −20 mV during the final 20 ms of stimulation.

In heatmaps of Figure 4, the explored parameter ranges were uniformly sampled at 10 equally spaced values, and the model was stimulated for 200 ms with an applied current equal to 20 *μ*A/cm^2^for the whole interval. Spike threshold was defined as the first point in the rising phase at which dV/dt > 10 mV/ms. Spike width was measured at half amplitude, where amplitude was defined as the voltage difference between the peak and threshold. The afterhyperpolarization amplitude was defined as the difference between the threshold voltage and the minimum voltage reached during the falling phase. Negative AHP values were retained as an extension of the classical definition and indicated the absence of afterhyperpolarization.

### Retrospective Nav1.7 mutation benchmarks and kinetic-sensitivity analysis

To evaluate the practical use of the reformulation for biologically reported channel perturbations, we modeled the Nav1.7-A863P mutation characterized by Harty et al^34^. The published voltage-clamp data report a hyperpolarizing shift in sodium activation from −15.9 mV in WT to −23.6 mV in A863P and a depolarizing shift in fast inactivation from −76.9 mV in WT to −67.1 mV in A863P. The accompanying current-clamp recordings in DRG neurons report depolarization of resting membrane potential from −56.9 to −50.4 mV, reduction of current threshold from 248 to 157 pA, increased firing frequency over suprathreshold current injections, and no significant change in spike amplitude^34^.

The mutation benchmarks evaluated the mapping of limited steady-state gating information onto excitability, rather than reconstructing a DRG neuron. WT and mutant conditions were constructed separately for each study so that each mutant was compared with its study-specific WT. No mutation-specific potassium measurements were used. The classical potassium gate was held identical between WT and A863P in the classical fitting analyses; the voltage-shift-only comparator used the additional common potassium shift specified below. In Hill benchmarks the potassium gate was specified by design and held identical between wild type and mutant within each case study. Its midpoint and slope were chosen so that potassium activation is recruited above the sodium activation range, preserving the classical ordering in which the delayed rectifier opposes depolarization only after the sodium-driven upstroke. The Hill parameters used were (− 110, 6.333, −15) for F1449V and (−130, 7, −21) for A1632G, with the classical βn retained; these values are listed in Supplementary Table 10. Neither formulation included Nav1.8 or the other conductances needed for a detailed DRG model. Agreement in the direction of an excitability change was evaluated separately from agreement in its magnitude. The comparison framework is summarized in Figure 5a, and the A863P implementations and their assumptions are collated in Supplementary Table 7. Steady-state targets and kinetic assumptions were specified separately. For a gate x, *x*_∞_(*V*) = (*α_x_*(*V*))/(*α_x_*(*V*) + *β_x_*(*V*)), whereas *τ_x_*(*V*) = 1/(*α_x_*(*V*) + *β_x_*(*V*)). A steady-state curve constrains the ratio of the two rates, but not their common scale.

Consequently, retaining a complementary rate, penalizing deviations from a reference time constant, and selecting a particular inactivation-rate parameterization are kinetic assumptions, even when no kinetic data are fitted. The Hill and classical comparisons were therefore organized by target curve and kinetic rule; not all implementations used identical inactivation targets. Classical HH rate equations were evaluated at *V* + 40 mV in place of *V*, and the same offset was applied to retained classical expressions in the modified model. This offset was part of the case-study implementation.

### Experimental gating inputs and curve reconstruction

For A863P, activation targets used WT *V*½ = −15.9 mV and *k* = 7.3 mV, and mutant *V*½ = −23.6 mV and *k* = 8.4 mV. Inactivation targets used WT *V*½ = −76.9 mV and *k* = 6.9 mV, and mutant *V*½ = −67.1 mV and *k* = 6.4 mV. Increasing activation targets were constructed as 1/1 + *exp*[−(*V* - *V*½)/*k*], and decreasing inactivation targets as 1/1 + *exp*[(*V* - *V*½)/*k*]. The classical A863P fits sampled these targets at 25 voltages. The experimental gating descriptors and the corresponding direct-mapping Hill parameters are listed in Supplementary Table 8.

The additional Hill-model benchmarks used gating descriptors for F1449V^51^ and A1632G^33^. For F1449V, activation (*V*½, *k*), in mV, was (−15.2, 7.65) for WT and (−22.8, 8.08) for mutant; inactivation was (−71.3, 6.68) and (−67.0, 7.00), respectively. For A1632G, activation was (−20.68, 8.09) for WT and (−27.17, 8.64) for mutant; inactivation was (−86.32, 7.58) and (−76.91, 6.75), respectively. In both studies the half-activation and half-inactivation voltages are reported and were used directly. The slope factors are not reported numerically. Unlike the reported A863P slopes, these were recovered from the published steady-state curves of Fig. 4A of ^52^ and Fig. 2C,D of ^51^ by selecting a non-midpoint curve value and solving he corresponding Boltzmann equation for *k*. They were treated as reconstructed descriptors rather than directly reported measurements. These inputs are collated in Supplementary Table 8; the resulting gating curves are shown in Supplementary Figure 5a,d.

### Direct Hill mapping and parameter specification

For direct mapping, sodium activation was represented by the Hill function *m*_∞_. The Hill midpoint was set equal to the reported activation *V*½. With the shift a fixed, matching the derivative at the midpoint gives l = (*V*½. −*a*)/*k*; thus, the shift must be specified in addition to the experimental midpoint and slope. The retained classical *β_m_* determined *α_m_* through *α_m_* = *β_m_*. *m*_∞_/(1 - *m*_∞_), and therefore also determined activation kinetics. No kinetic measurements were fitted in this direct implementation.

The activation Hill parameter triples (*a*, *V*½, l), with voltages in mV, were: A863P WT (−250, - 15.9, 32.0685) and mutant (−250, −23.6, 26.9524); F1449V WT (−250, −15.2, 30.6760) and mutant (−250, −22.8, 28.1319); and A1632G WT (−250, −20.68, 28.3461) and mutant (−250, −27.17, 25.7905). These parameter sets are tabulated in Supplementary Table 8.

For inactivation, the Hill function described *β*_ℎ_, not ℎ_∞_. Classical *α*_ℎ_ was retained, so ℎ_∞_ = *α*_ℎ_/(*α*_ℎ_ + *β*_ℎ_) is determined by the ratio of the two rates. The reported half-inactivation voltage therefore fixes one condition on that ratio: the two rates must be equal at the published midpoint. Once the shift and the Hill exponent are chosen, this condition determines the Hill midpoint of *β*_ℎ_ uniquely, so the midpoint was imposed rather than tuned.

The *β*_ℎ_ Hill midpoint is therefore not the experimental half-inactivation voltage. The *β*_ℎ_ parameter triples (*a*, *V*½, l) were: A863P WT (−150, −73, 4.88) and mutant (−150, −47, 4.68); F1449V WT (−150, −57.5, 4.4) and mutant (−150, −39, 3.5); and A1632G WT (−130, −96.576, 3.4) and mutant (−120, −70.024, 2).

The remaining parameters *a* and l set the steepness of the derived ℎ∞. These were selected to follow the direction of the reported slope change between WT and mutant rather than to reproduce its magnitude: a larger reported shift was represented by a larger shift in the Hill midpoint, and a steeper reported curve by a steeper βh. With this rule the derived ℎ∞ reproduces the published half-inactivation voltage to within 0.5 mV in all six conditions (Supplementary Table 9), and reproduces the direction of the slope change in all three mutations. This construction does not establish that the full experimental inactivation slope or time course is reproduced. The inactivation descriptors and rate-function parameters are distinguished in Supplementary Table 8.

### A863P Hill ensembles

Two complementary Hill ensembles assessed sensitivity to the fitting initialization and to the retained inactivation rate. In each, Hill parameters were refitted from 200 random starting points within fixed bounds, where each start fitted the WT and mutant curves together as one paired set, so that every reported mutant/WT ratio comes from a single paired fit. The activation target was the Boltzmann *m*_∞_ curve constructed from the reported midpoint and slope. The inactivation target was the ℎ_∞_ curve chosen in the direct Hill implementation, rather than the published Boltzmann inactivation curve. Fits were accepted when the root-mean-square error (RMSE) of every steady-state fit was at most 0.02 in dimensionless gating units. Thus, these ensembles tested reproducibility conditional on a selected inactivation target. Their excitability summaries are included in Figure 5b and Supplementary Table 7. Both steady-state targets were evaluated on fixed 25-point voltage grids spanning the range plotted in the source study: −80 to 40 mV for *m*_∞_, −130 to 10 mV for ℎ_∞_ evenly spaced. Starting points were drawn independently for every fit within the fixed bounds, log-uniformly for the strictly positive constants and uniformly for the voltage constants, with the random number generator seeded once per ensemble (seed 1) before the draw loop, so that each ensemble is reproducible as a complete sequence. Each fit minimised the residual between the model steady state and its target by bounded nonlinear least squares (MATLAB lsqnonlin) with function and step tolerances of 10⁻¹⁴, at most 3 × 10⁴ function evaluations and 3 × 10³ iterations; these tolerances constitute the stopping criteria, and no constraints were imposed beyond the bounds. Wild-type and mutant were fitted within the same draw, each from its own independently drawn starting point, and a draw was retained only if all four steady-state fits of that draw met the acceptance criterion.

In the fixed-complementary-rate ensemble, *β_m_* and *α*_ℎ_ retained their classical forms. In the second ensemble, the amplitude and voltage-scale parameters of exponential *α*_ℎ_ were fitted jointly with the Hill *β*_ℎ_ parameters. The free parameters were (*a*, l, *V*½) for activation and (*a*, l, *V*½, *D*_ℎ_, *E*_ℎ_) for inactivation, bounded below by (−400, 2, −50) and (−300, 1, −110, 0.014, 4) and above by (−250, 80, 10) and (−150, 60, 10, 0.35, 100) respectively, with the same bounds used for every draw and are summarized is supplementary Table 14. Classical *β_m_* was still retained because the activation steady-state target alone did not identify its kinetics. This retention was an explicit modeling choice, rather than a uniquely determined consequence of the steady-state measurements. For both ensembles, the resulting WT and mutant excitability metrics were evaluated without fitting current-clamp outputs. The free-*α*_ℎ_ ensemble rheobase ratios are shown in Supplementary Figure 6b, with the associated Hill-model time constants in Supplementary Figure 4e-h.

### Classical HH sensitivity comparisons for A863P

All-rate-parameter fits used the classical sodium rate-equation forms, with all five parameters per gate fitted jointly across alpha and beta to the WT and A863P Boltzmann targets by nonlinear least squares. No kinetic penalty was applied. Fits were repeated from 200 random starting points within fixed bounds; a solution was accepted only when every steady-state fit had RMSE at most 0.02. Bounds were fixed across draws and centered on the classical constants translated along the voltage axis so that the gate midpoint coincided with the target midpoint; for the multiplicative constants they spanned one fifth to five times the center value, and for the voltage constant of each gate ±30 mV about the center. Starting points, optimizer settings, stopping criteria, acceptance and the wild-type/mutant pairing rule were as described for the Hill ensembles above.

In the fixed-complementary-rate variant, *β_m_* and *α*_ℎ_ were fixed to their classical forms and only the remaining rate parameters were fitted, again using 200 starts and the same acceptance criterion. These alternatives distinguish non-uniqueness of steady-state fits from the consequences of fixing a particular rate. The implementation comparison is shown in Figure 5b and Supplementary Table 7; the all-free classical rheobase-ratio ensemble is shown in Supplementary Figure 6a, and classical ensemble time constants in Supplementary Figure 4a-d.

The voltage-shift-only comparator evaluated both rates of each sodium gate at *V* - *ΔV*, choosing *ΔV* so that the resulting steady-state midpoint equalled the study-specific WT or A863P midpoint. No slope fitting was performed: the classical steady-state slope and the time-constant shape were retained under voltage translation. The potassium *n*_∞_ relationship received the same −0.88 mV WT activation shift in both conditions. This comparator therefore used midpoint information only and was not equivalent to fitting the full Boltzmann targets. This comparator is included in Figure 5b and Supplementary Table 7.

A separate consistency check fitted classical HH rates, with complementary rates fixed, to published Boltzmann activation targets and to the ℎ_∞_ curve generated by the direct Hill implementation. This comparison reproduced a model-derived inactivation target and was not treated as an independent biological benchmark or as evidence that the published inactivation curve had been fitted equally by both formulations. The detailed comparison using the Hill-derived inactivation target is shown in Supplementary Figure 3, with the consistency-check summary in Figure 5b and Supplementary Table 7.

### Time-constant constraints and rescaling

For time-constant-constrained classical fits, all five sodium rate parameters per gate were fitted to the Boltzmann steady-state targets while penalizing deviation from a reference *τ*(*V*). The reference was the classical HH time constant translated by the voltage difference between its classical steady-state midpoint and the reported midpoint for the corresponding gate and WT or mutant condition. The objective was *L* = ∑_i_[*x* _∞(*Vi*)_ - *x*(*target*)(*V_i_*)]^2^ + *λ*^2^ ∑_i_[log *τ*(*V*_i_) - log *τ*_(*target*)(*Vi*)_]^2^. The logarithmic penalty constrained relative rather than absolute kinetic deviations. Fits used lambda = 0.02, 0.05, and 0.1, with 0.05 designated as the reference. These reference kinetics were assumed rather than experimentally measured mutation-specific kinetics. These fits used their own protocol rather than that of the unconstrained ensembles. Targets were Boltzmann curves evaluated analytically from the published descriptors on 25-point grids. Parameter bounds were fixed absolute intervals rather than intervals centered on the classical constants: (10⁻³, −200, 1, 10⁻³, 2) to (50, 300, 60, 100, 200) for the activation gate and (10⁻⁵, 2, −200, 1, 10⁻³) to (10, 200, 300, 60, 50) for the inactivation gate, these boundaries are summarized in Supplementary Table 15. Each gate and condition was fitted from a fixed set of seven starting points, obtained by perturbing the voltage-shifted classical parameter vector, and the solution with the lowest total residual was retained; these starting points are identical on every run and no random multistart was used. Because these fits trace the effect of the constraint weight rather than sample an ensemble, no RMSE acceptance threshold was applied and every fit is reported; fit quality is characterized instead by the maximum deviation from the steady-state target and by the range of the ratio between the fitted and reference time constants. The optimizer and its tolerances were as described above. The corresponding excitability comparisons are summarized in Figure 5b and Supplementary Table 7.

Starting from the *λ* = 0.05 reference fit, both rates of the activation gate were multiplied by ∅*_m_* and both rates of the inactivation gate by ∅_ℎ_. Each factor took values {0.25, 0.5, 1, 2, 4}, giving 25 combinations. The same gate-specific factors were applied to WT and mutant. Multiplying both rates by phi leaves *x*_∞_ unchanged and divides *τ* by ∅, isolating kinetic sensitivity while preserving the fitted steady-state curves. Random-start ensembles and rescaling grids were treated as sensitivity analyses, not as biological replicates or probability distributions over mutation outcomes. The two-factor kinetic-rescaling grid is shown in Figure 5c, with its rheobase-ratio summary in Figure 5b and Supplementary Table 7.

Rheobase was estimated as described above, by a coarse scan followed by bisection to within 0.05 µ*A*/*cm*², identifying the smallest current that elicited at least one detected spike during the stimulation interval. Activation-only, inactivation-only, and combined perturbations were simulated separately. Frequency-current curves for the A863P benchmark were generated by sweeping *I_app_* over 0-140 *μA*/*cm*² and calculating firing rate from the number of detected spikes during the stimulation interval. First-spike features were extracted using the same spike-detection, threshold, width, amplitude, and afterhyperpolarization definitions described above.

### Mutation-benchmark readouts and experimental comparison

For A863P sensitivity analyses, a 100-ms current step was applied from 50 to 150 ms within a 200-ms simulation. Simulations started from an initial membrane potential of −50 mV with the gating variables initialized as described before, were integrated with MATLAB’s ode113 solver at relative and absolute tolerances of 10⁻⁶and 10⁻⁸,and spikes were detected using the amplitude-based rule described above. Rheobase was taken as the smallest step amplitude eliciting at least one detected action potential during the stimulus. It was located by a coarse scan over 0 - 200 µ*A*/*cm*², followed by bisection of the bracketing interval; the coarse scan used 2 µ*A*/*cm*²increments with bisection continued until the interval was narrower than 0.05µ*A*/*cm*². The primary endpoint was the mutant/WT rheobase ratio; spike counts at 40 µ*A*/*cm*² were a secondary endpoint. Firing rates were computed from spikes during the 100-ms stimulation interval divided by 0.1 s. Activation-only, inactivation-only, and combined perturbations distinguished the contributions of the two gating changes in the direct Hill benchmarks. The F1449V and A1632G direct benchmarks used the same stimulus protocol, initial conditions, spike detection and rheobase search, and additionally included responses and first-spike features at 40 µ*A*/*cm*², using the feature definitions introduced before, and frequency–current comparisons obtained by sweeping *I_app_* from 0 to 140 µ*A*/*cm*². Rheobase contributions from activation-only, inactivation-only, and combined direct Hill perturbations are reported in Supplementary Table 11. F1449V and A1632G voltage responses and frequency-current curves are shown in Supplementary Figure 5b,c,e,f, and their first-spike features are listed in Supplementary Table 13.

Experimental reference ratios were calculated within each study from mutant and WT current thresholds: 157/248 = 0.633 for A863P, 93.1/124.1 = 0.750 for F1449V, and 102/158 = 0.646 for A1632G. These dimensionless ratios allowed comparison with model current densities without converting experimental pA values to µ*A*/*cm*². A ratio below one indicated the reported direction of threshold reduction; its distance from the experimental ratio described quantitative discrepancy. Experimental current-clamp measurements were used for retrospective comparison, not as fitting targets. Differences in cellular conductances, stimulus protocols, and experimental preparations limit direct comparison of absolute currents, firing rates, and waveform measurements. The additional F1449V/A1632G analyses concerned the Hill model; the systematic classical HH kinetic-sensitivity comparisons concerned A863P. The comparison of experimental current-threshold ratios with direct Hill-model rheobase ratios is presented in Figure 5d and Supplementary Table 12.

## Data availability

The historical Hodgkin-Huxley data used for model fitting were obtained from the original published studies cited in the manuscript. All data generated through numerical simulations and used to produce the figures and analyses in this study are available within the article and its Supplementary Information. Additional processed data files are available from the corresponding author upon reasonable request.

## Code availability

The MATLAB code used to implement the classical and Hill-based Hodgkin-Huxley models, perform sigmoidal fitting, generate excitability maps, extract spike features, and reproduce the figures will be made publicly available through https://github.com/daoulab at a specified repository. The archived version corresponding to the published manuscript will include the model equations, fitted parameters, simulation scripts, and figure-generation code.

## Author contributions

B.S. contributed to model implementation, simulations, formal analysis, visualization, and preparation of the original manuscript draft. J.F. contributed to mathematical analysis, validation, interpretation of the model formulation, and critical revision of the manuscript. A.D. conceived and supervised the study, guided the interpretation of the results, administered the project, and prepare and revised the manuscript. All authors reviewed and approved the final manuscript.

## References

1. Hille, B. Ion Channels of Excitable Membranes. (Sinauer, Sunderland, Mass, 2001).

2. Hodgkin, A. L. & Huxley, A. F. A quantitative description of membrane current and its application to conduction and excitation in nerve. The Journal of Physiology 117, 500–544 (1952).

3. Hodgkin, A. L. & Huxley, A. F. The dual effect of membrane potential on sodium conductance in the giant axon of *Loligo*. The Journal of Physiology 116, 497–506 (1952).

4. Hodgkin, A. L. & Huxley, A. F. The components of membrane conductance in the giant axon of *Loligo*. The Journal of Physiology 116, 473–496 (1952).

5. Hodgkin, A. L. & Huxley, A. F. Currents carried by sodium and potassium ions through the membrane of the giant axon of *Loligo*. The Journal of Physiology 116, 449–472 (1952).

6. Hodgkin, A. L., Huxley, A. F. & Katz, B. Measurement of current-voltage relations in the membrane of the giant axon of *Loligo*. The Journal of Physiology 116, 424–448 (1952).

7. FitzHugh, R. Impulses and Physiological States in Theoretical Models of Nerve Membrane. Biophysical Journal 1, 445–466 (1961).

8. John Rinzel & G. Bard Ermentrout. Analysis of neural excitability and oscillations. in Methods in Neuronal Modeling: From Synapses to Networks (eds Christof Koch & Idan Segev) 135–169 (MIT Press, 1989).

9. Morris, C. & Lecar, H. Voltage oscillations in the barnacle giant muscle fiber. Biophysical Journal 35, 193–213 (1981).

10. Nagumo, J., Arimoto, S. & Yoshizawa, S. An Active Pulse Transmission Line Simulating Nerve Axon. Proc. IRE 50, 2061–2070 (1962).

11. Izhikevich, E. M. Simple model of spiking neurons. IEEE Trans. Neural Netw. 14, 1569–1572 (2003).

12. Izhikevich, E. M. Dynamical Systems in Neuroscience: The Geometry of Excitability and Bursting. (The MIT Press, 2006). doi:10.7551/mitpress/2526.001.0001.

13. Bhalla, U. S. & Bower, J. M. Exploring parameter space in detailed single neuron models: simulations of the mitral and granule cells of the olfactory bulb. Journal of Neurophysiology 69, 1948–1965 (1993).

14. Bekkers, J. M. Distribution and activation of voltage-gated potassium channels in cell-attached and outside-out patches from large layer 5 cortical pyramidal neurons of the rat. The Journal of Physiology 525, 611–620 (2000).

15. Destexhe, A. & Huguenard, J. R. Nonlinear Thermodynamic Models of Voltage-Dependent Currents. J Comput Neurosci 9, 259–270 (2000).

16. Bower, J. M. & Beeman, D. The Book of GENESIS. (Springer New York, New York, NY, 1998). doi:10.1007/978-1-4612-1634-6.

17. Hines, M. L. & Carnevale, N. T. The NEURON Simulation Environment. Neural Computation 9, 1179–1209 (1997).

18. Krouchev, N. I., Rattay, F., Sawan, M. & Vinet, A. From Squid to Mammals with the HH Model through the Nav Channels’ Half-Activation-Voltage Parameter. PLoS ONE 10, e0143570 (2015).

19. Alon, U. An Introduction to Systems Biology: Design Principles of Biological Circuits. (Chapman and Hall/CRC, 2006). doi:10.1201/9781420011432.

20. A.V., H. The possible effects of the aggregation of the molecules of haemoglobin on its dissociation curves. Journal of Physiology 40, iv–vii (1910).

21. Santillán, M. On the Use of the Hill Functions in Mathematical Models of Gene Regulatory Networks. Math. Model. Nat. Phenom. 3, 85–97 (2008).

22. Weiss, J. N. The Hill equation revisited: uses and misuses. The FASEB Journal 11, 835–841 (1997).

23. Bezanilla, F. The Voltage Sensor in Voltage-Dependent Ion Channels. Physiological Reviews 80, 555–592 (2000).

24. Chowdhury, S. & Chanda, B. Estimating the voltage-dependent free energy change of ion channels using the median voltage for activation. Journal of General Physiology 139, 3–17 (2012).

25. Gesztelyi, R. et al. The Hill equation and the origin of quantitative pharmacology. Arch. Hist. Exact Sci. 66, 427–438 (2012).

26. Sigworth, F. J. Voltage gating of ion channels. Quart. Rev. Biophys. 27, 1–40 (1994).

27. Bezanilla, F. How membrane proteins sense voltage. Nat Rev Mol Cell Biol 9, 323–332 (2008).

28. Colquhoun, D. & Hawkes, A. G. The Principles of the Stochastic Interpretation of Ion-Channel Mechanisms. in Single-Channel Recording (eds Sakmann, B. & Neher, E.) 397–482 (Springer US, Boston, MA, 1995). doi:10.1007/978-1-4419-1229-9_18.

29. Yellen, G. The moving parts of voltage-gated ion channels. Quart. Rev. Biophys. 31, 239–295 (1998).

30. Achard, P. & De Schutter, E. Complex Parameter Landscape for a Complex Neuron Model. PLoS Comput Biol 2, e94 (2006).

31. Marder, E. & Goaillard, J.-M. Variability, compensation and homeostasis in neuron and network function. Nat Rev Neurosci 7, 563–574 (2006).

32. Prinz, A. A., Bucher, D. & Marder, E. Similar network activity from disparate circuit parameters. Nat Neurosci 7, 1345–1352 (2004).

33. Van Geit, W., De Schutter, E. & Achard, P. Automated neuron model optimization techniques: a review. Biol Cybern 99, 241–251 (2008).

34. Harty, T. P. et al. Nav 1.7 Mutant A863P in Erythromelalgia: Effects of Altered Activation and Steady-State Inactivation on Excitability of Nociceptive Dorsal Root Ganglion Neurons. J. Neurosci. 26, 12566–12575 (2006).

35. Strogatz, S. H. Nonlinear Dynamics and Chaos: With Applications to Physics, Biology, Chemistry, and Engineering. (Addison-Wesley publ, Reading (Mass.), 1994).

36. Ermentrout, G. B. & Terman, D. H. Mathematical Foundations of Neuroscience. vol. 35 (Springer New York, New York, NY, 2010).

37. Rinzel, J. & Miller, R. N. Numerical calculation of stable and unstable periodic solutions to the Hodgkin-Huxley equations. Mathematical Biosciences 49, 27–59 (1980).

38. Kuznetsov, Y. A. Elements of Applied Bifurcation Theory. vol. 112 (Springer New York, New York, NY, 2004).

39. Goldman, M. S., Golowasch, J., Marder, E. & Abbott, L. F. Global Structure, Robustness, and Modulation of Neuronal Models. J. Neurosci. 21, 5229–5238 (2001).

40. Armstrong, C. M. Sodium channels and gating currents. Physiological Reviews 61, 644–683 (1981).

41. Bean, B. P. The action potential in mammalian central neurons. Nat Rev Neurosci 8, 451–465 (2007).

42. Platkiewicz, J. & Brette, R. A Threshold Equation for Action Potential Initiation. PLoS Comput Biol 6, e1000850 (2010).

43. Bhatt, D. H., Zhang, S. & Gan, W.-B. Dendritic Spine Dynamics. Annu. Rev. Physiol. 71, 261–282 (2009).

44. Călin, A., Ilie, A. S. & Akerman, C. J. Disrupting Epileptiform Activity by Preventing Parvalbumin Interneuron Depolarization Block. J. Neurosci. 41, 9452–9465 (2021).

45. Izhikevich, E. M. NEURAL EXCITABILITY, SPIKING AND BURSTING. Int. J. Bifurcation Chaos 10, 1171–1266 (2000).

46. Prescott, S. A., De Koninck, Y. & Sejnowski, T. J. Biophysical Basis for Three Distinct Dynamical Mechanisms of Action Potential Initiation. PLoS Comput Biol 4, e1000198 (2008).

47. Azouz, R. & Gray, C. M. Dynamic spike threshold reveals a mechanism for synaptic coincidence detection in cortical neurons *in vivo*. Proc. Natl. Acad. Sci. U.S.A. 97, 8110–8115 (2000).

48. Naundorf, B., Wolf, F. & Volgushev, M. Unique features of action potential initiation in cortical neurons. Nature 440, 1060–1063 (2006).

49. Mainen, Z. F. & Sejnowski, T. J. Influence of dendritic structure on firing pattern in model neocortical neurons. Nature 382, 363–366 (1996).

50. McCormick, D. A., Connors, B. W., Lighthall, J. W. & Prince, D. A. Comparative electrophysiology of pyramidal and sparsely spiny stellate neurons of the neocortex. Journal of Neurophysiology 54, 782–806 (1985).

51. Dib-Hajj, S. D. et al. Gain-of-function mutation in Nav1.7 in familial erythromelalgia induces bursting of sensory neurons. Brain 128, 1847–1854 (2005).

52. Yang, Y. et al. Nav1.7-A1632G Mutation from a Family with Inherited Erythromelalgia: Enhanced Firing of Dorsal Root Ganglia Neurons Evoked by Thermal Stimuli. Journal of Neuroscience 36, 7511–7522 (2016).

53. Borg-Graham, L. J. Modelling the non-linear conductances of excitable membranes. in Cellular Neurobiology (eds Chad, J. & Wheal, H.) 247–275 (Oxford University PressOxford, 1991). doi:10.1093/oso/9780199631063.003.0013.

54. Brette, R. & Gerstner, W. Adaptive Exponential Integrate-and-Fire Model as an Effective Description of Neuronal Activity. Journal of Neurophysiology 94, 3637–3642 (2005).

55. Model Selection and Multimodel Inference. (Springer New York, New York, NY, 2002). doi:10.1007/b97636.

56. Akaike, H. A new look at the statistical model identification. IEEE Trans. Automat. Contr. 19, 716–723 (1974).

57. Anderson, T. W. & Darling, D. A. Asymptotic Theory of Certain ‘Goodness of Fit’ Criteria Based on Stochastic Processes. Ann. Math. Statist. 23, 193–212 (1952).

58. Colquhoun, D. & Hawkes, A. G. On the stochastic properties of single ion channels. Proceedings of the Royal Society of London. Series B. Biological Sciences 211, 205–235 (1981).

59. Fink, M. & Noble, D. Markov models for ion channels: versatility versus identifiability and speed. Phil. Trans. R. Soc. A. 367, 2161–2179 (2009).

60. Vandenberg, C. A. & Bezanilla, F. A sodium channel gating model based on single channel, macroscopic ionic, and gating currents in the squid giant axon. Biophysical Journal 60, 1511–1533 (1991).

61. Single-Channel Recording. (Springer US, Boston, MA, 1995). doi:10.1007/978-1-4419-1229-9.

62. Frankenhaeuser, B. & Hodgkin, A. L. The action of calcium on the electrical properties of squid axons. The Journal of Physiology 137, 218–244 (1957).

63. Huxley, A. F. ION MOVEMENTS DURING NERVE ACTIVITY. Annals of the New York Academy of Sciences 81, 221–246 (1959).

64. Sterratt, D. C. Q10: The Effect of Temperature on Ion Channel Kinetics. in Encyclopedia of Computational Neuroscience (eds Jaeger, D. & Jung, R.) 2949–2950 (Springer New York, New York, NY, 2022). doi:10.1007/978-1-0716-1006-0_236.

65. Dayan, P. & Abbott, L. F. Theoretical Neuroscience: Computational and Mathematical Modeling of Neural Systems. (Massachusetts Institute of Technology Press, Cambridge, Mass, 2001).

66. Keener, J. & Sneyd, J. Mathematical Physiology. vol. 8 (Springer New York, New York, NY, 1998).

67. Hodgkin, A. L. & Katz, B. The effect of temperature on the electrical activity of the giant axon of the squid. The Journal of Physiology 109, 240–249 (1949).

68. Traub, R. D. & Miles, R. Neuronal Networks of the Hippocampus. (Cambridge University Press, 1991). doi:10.1017/CBO9780511895401.

69. Wang, X.-J. & Buzsáki, G. Gamma Oscillation by Synaptic Inhibition in a Hippocampal Interneuronal Network Model. J. Neurosci. 16, 6402–6413 (1996).

70. Cole, K. S. Dynamic electrical characteristics of the squid axon membrane. Archives des Sciences Physiologiques 3, 253–258 (1949).

71. Marmont, G. Studies on the axon membrane. I. A new method. J. Cell. Comp. Physiol. 34, 351–382 (1949).

72. MATLAB Optimization Toolbox. MathWorks Inc. (2023).

73. Press, W. H., Teukolsky, S. A., William T. V. & Brian P. F. Numerical Recipes: The Art of Scientific Computing. (Cambridge University Press, Cambridge, UK; New York, 2007).

