## Supplementary Figures and Tables for "Hill-Based Reformulation of the Hodgkin-Huxley Model for Mapping Gating Changes to Neuronal Excitability"

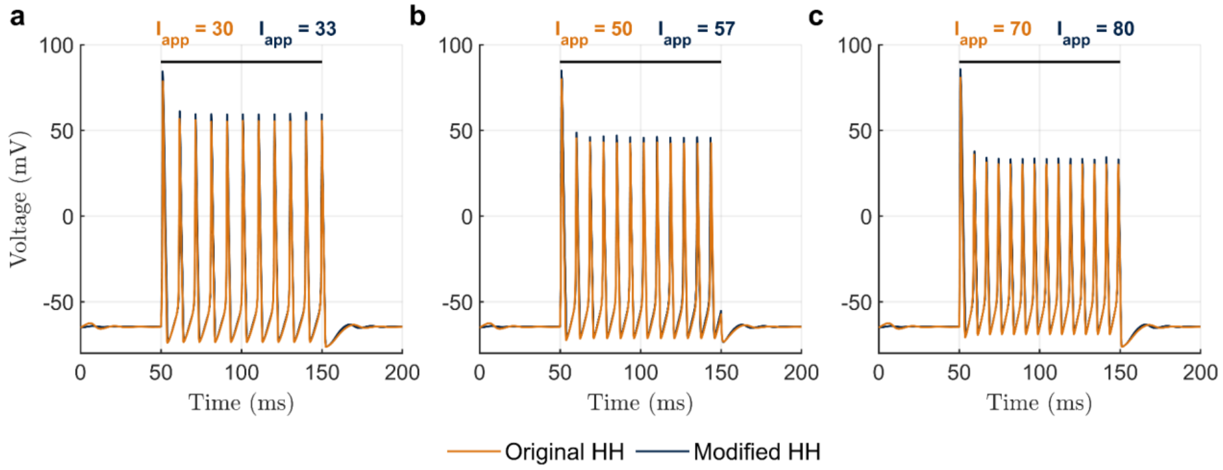


**Supplementary Figure 1. Representative voltage traces illustrating HH-like firing after current adjustment in the Hill-based model.** Membrane-potential traces generated by the original Hodgkin-Huxley model (HH; orange) and the Hill-based HH model (navy). **a-c**, Applied-current densities for the original and Hill-based models are 30 and 33, 50 and 57, and 70 and 80 μA/cm², respectively. Black bars indicate the 100-ms current steps applied from 50 to 150 ms. At these paired current levels, the models show closely matched spike timing and broadly similar waveforms, with visible differences in spike peaks and afterhyperpolarization. These examples illustrate preservation of repetitive firing after adjustment of applied current, consistent with the current-remapping and frequency-current analyses in Figure 2.

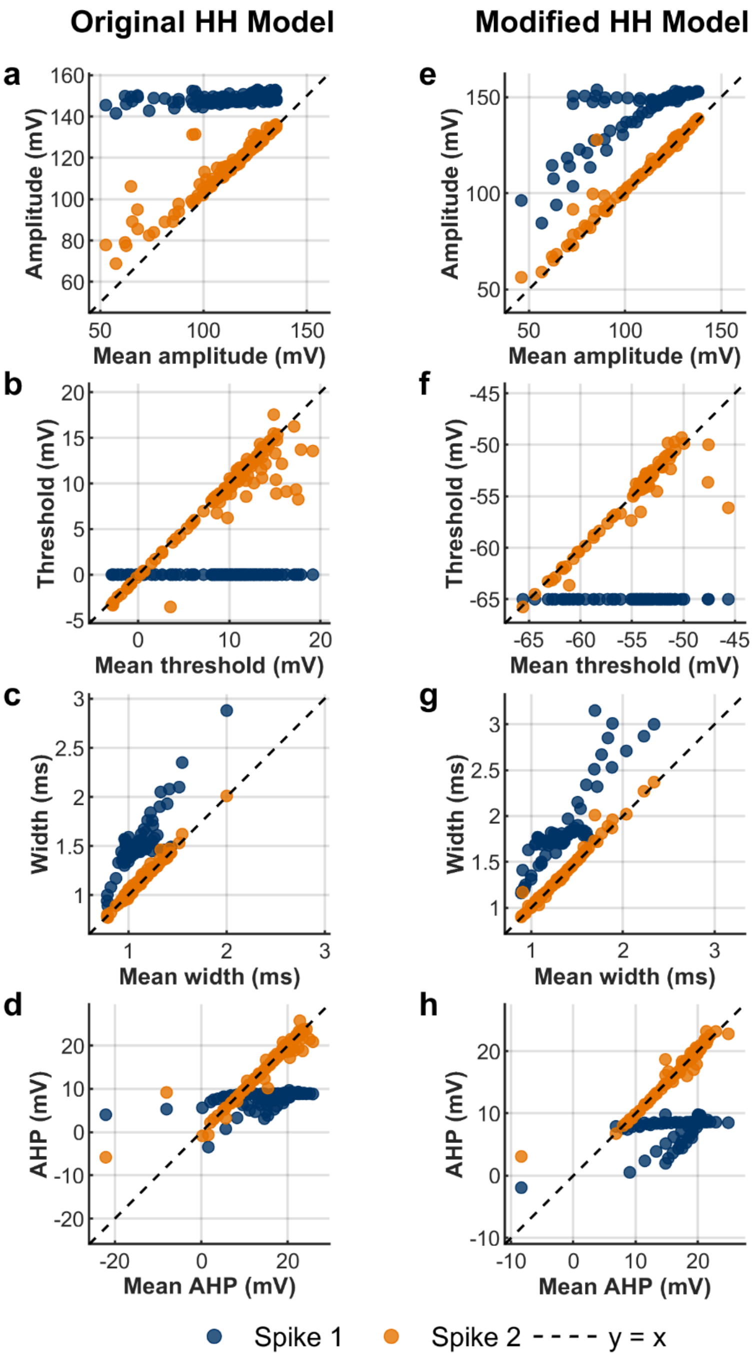


**Supplementary Figure 2. Comparison of first- and second-spike features with later spikes during repetitive firing.** Feature values for the first spike (blue) and second spike (orange) are plotted against the mean of the corresponding feature across spikes 3 to the end of the same train. **a-d**, Original Hodgkin-Huxley (HH) model. **e-h**, Hill-based HH model. Features are spike amplitude (**a**, **e**), voltage threshold (**b**, **f**), width (**c**, **g**), and afterhyperpolarization (AHP; **d**, **h**). Each point represents a simulated spike train; the x-axis gives the later-spike mean, and the y-axis gives the first- or second-spike value. Dashed black lines indicate identity (y = x). Across the sampled trains, second-spike measurements generally lie closer to the later-spike mean than first-spike measurements, supporting their use in the spike-feature sensitivity analysis in Figure 4.

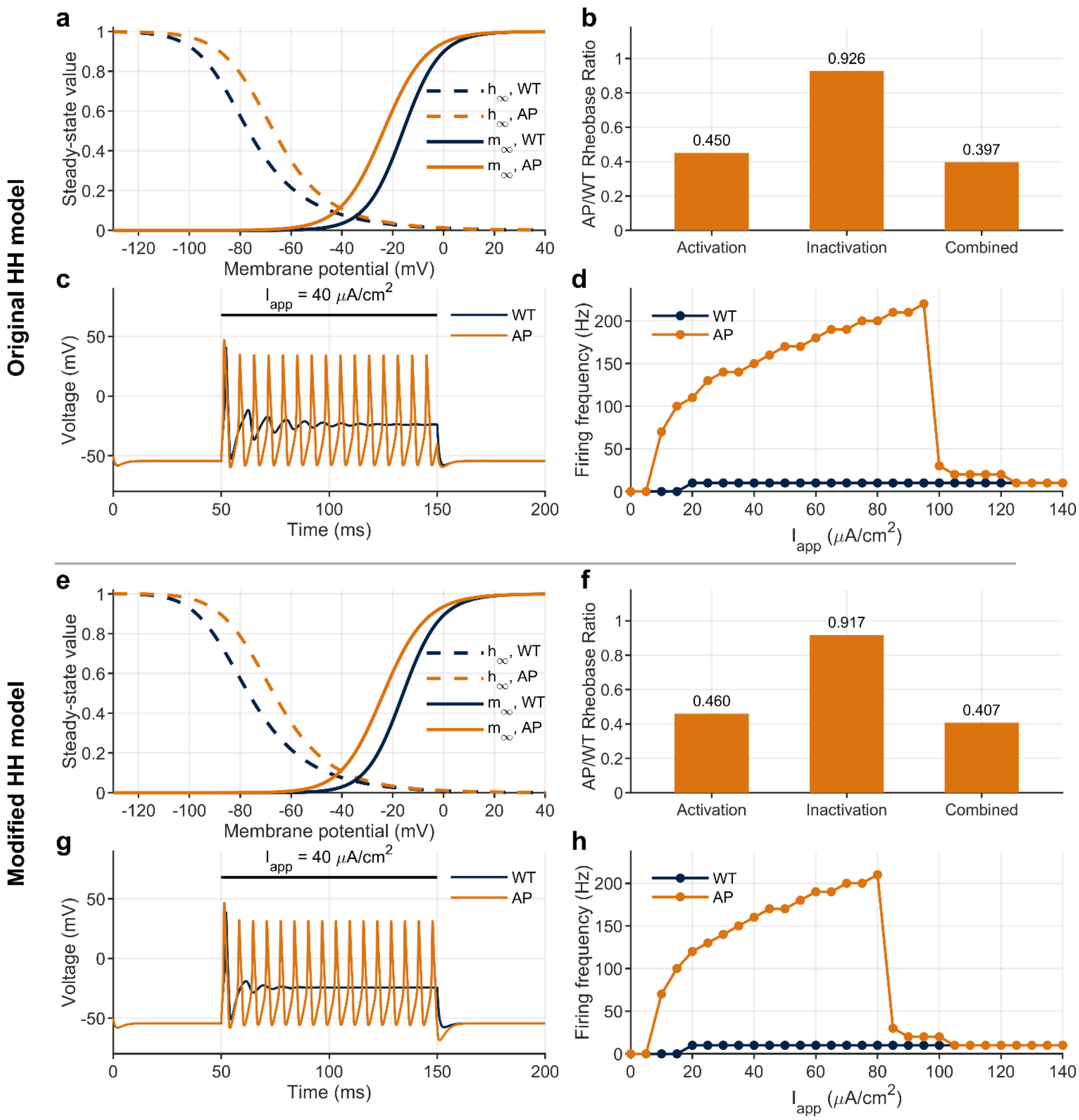


**Supplementary Figure 3. A863P fitting-based consistency check between classical and Hill-based HH implementations. a-d**, Classical Hodgkin-Huxley (HH) implementation fitted using Hill-derived inactivation as a target. **e-h**, Hill-based HH implementation. This comparison tests agreement between implementations under the selected gating construction rather than independent prediction from experimental steady-state descriptors alone. **a**, **e**, Steady-state sodium activation ($m_\infty$; solid lines) and inactivation ($h_\infty$; dashed lines) for wild type (WT; navy) and A863P (AP; orange). **b**, **f**, Mutant-to-WT rheobase ratios for activation-only, inactivation-only, and combined perturbations; values are indicated above the bars. **c**, **g**, Representative membrane-potential responses to an applied current of 40 μA/cm² from 50 to 150 ms. **d**, **h**, Firing frequency as a function of applied current. Both implementations reproduce reduced rheobase and repetitive mutant firing under these assumptions. Their agreement supports consistency between the selected implementations, rather than superior biological predictive accuracy of either formulation. Implementation details are summarized in Supplementary Table S7.

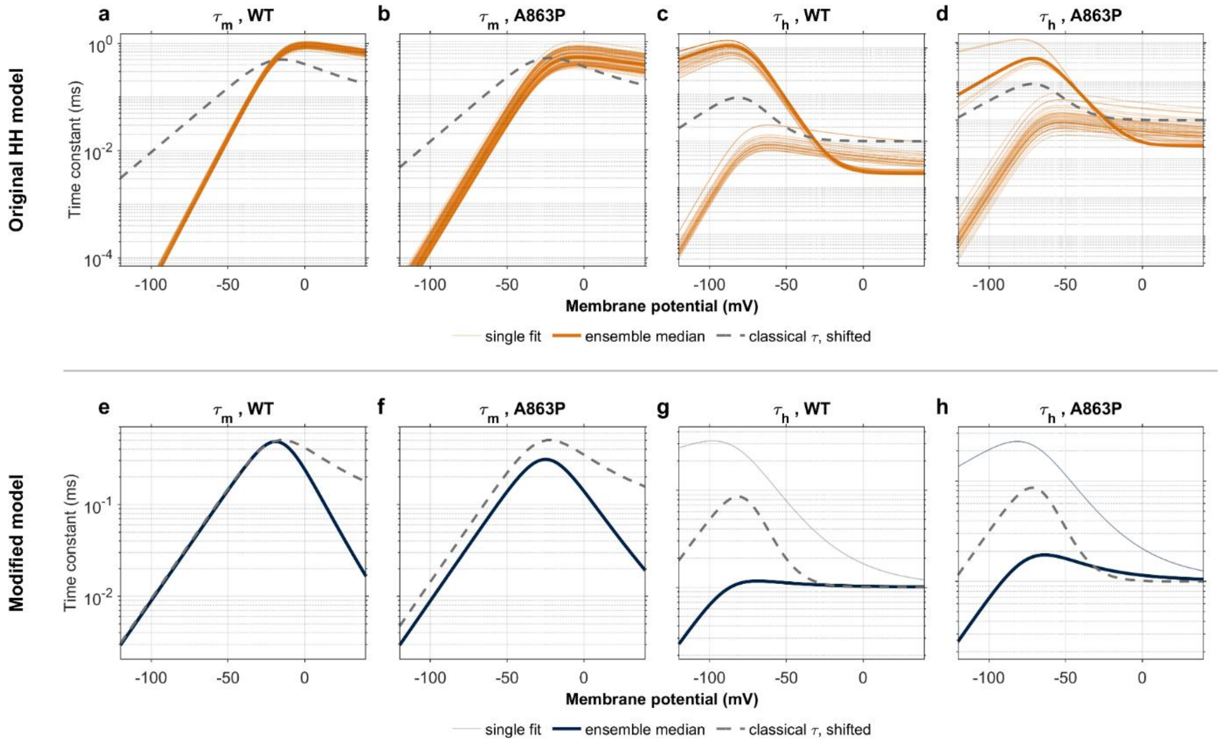


**Supplementary Figure 4. Sodium-gating time constants across A863P fitting ensembles.** Voltage dependence of the activation time constant, $\tau_m$, and inactivation time constant, $\tau_h$, for wild type (WT) and A863P in classical Hodgkin-Huxley (HH; **a-d**, orange) and Hill-based HH (**e-h**, navy) implementations. The classical panels show the ensemble with all rate parameters free, and the Hill panels show the ensemble in which the inactivation rate $\alpha_h$ is fitted jointly with the Hill $\beta_h$ parameters. Each panel draws 80 of the accepted fits. **a**, **e**, WT $\tau_m$; **b**, **f**, A863P $\tau_m$; **c**, **g**, WT $\tau_h$; **d**, **h**, A863P $\tau_h$. Thin lines show individual fits, thick lines indicate ensemble medians, and dashed gray lines show the voltage-shifted classical HH time constants used as references. Time constants are calculated from the forward and backward gating rates as $\tau_x$ = $1/(\tau_x + \beta_x)$. Time-constant axes are logarithmic, with different ranges between rows. The ensembles illustrate the kinetic variability compatible with the selected steady-state targets and rate assumptions. Classical and Hill-based ensembles use different inactivation targets and therefore do not constitute a comparison under identical constraints (Supplementary Table S7).

.

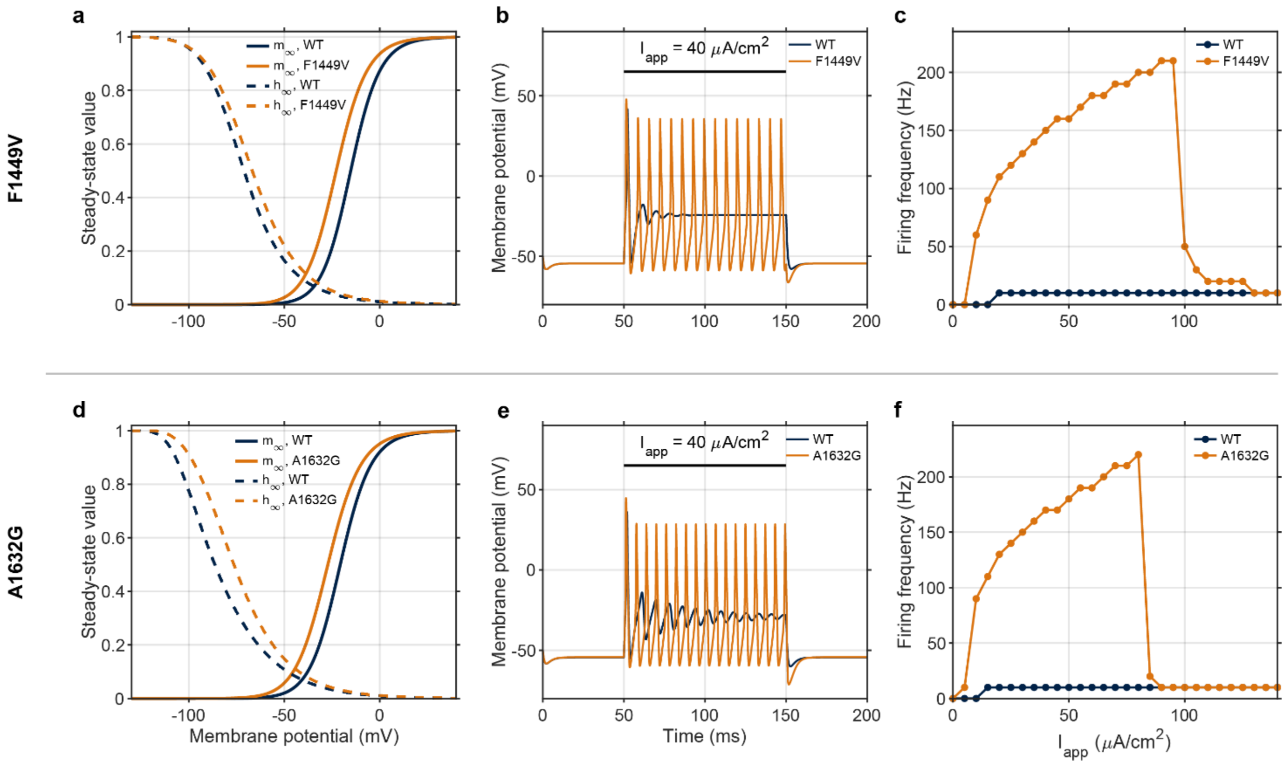


**Supplementary Figure 5. Hill-based predictions of gating and firing changes for the Nav1.7 mutations F1449V and A1632G.** Direct low-information mappings for F1449V (**a-c**) and A1632G (**d-f**), each compared with its study-specific wild-type (WT) control. WT is shown in navy and mutant in orange. **a**, **d**, Steady-state sodium activation ($m_\infty$; solid lines) and inactivation ($h_\infty$; dashed lines) resulting from the selected gating descriptors and model assumptions. **b**, **e**, Membrane-potential responses to an applied current density of 40 μA/cm² from 50 to 150 ms. Both mutants exhibit repetitive firing, whereas their corresponding WT models produce a single spike. **c**, **f**, Firing frequency as a function of applied current density. Mutant models show firing at lower currents and higher discharge rates over the repetitive-firing range, followed by a sharp decline at stronger drive. These simulations extend the qualitative mutation benchmarks to F1449V and A1632G under the specified Hill-based mapping; they do not compare predictive performance with classical HH. Gating descriptors and Hill parameters, rheobases, experimental threshold comparisons, and first-spike features are provided in Supplementary Tables S8, S11, S12, and S13, respectively.

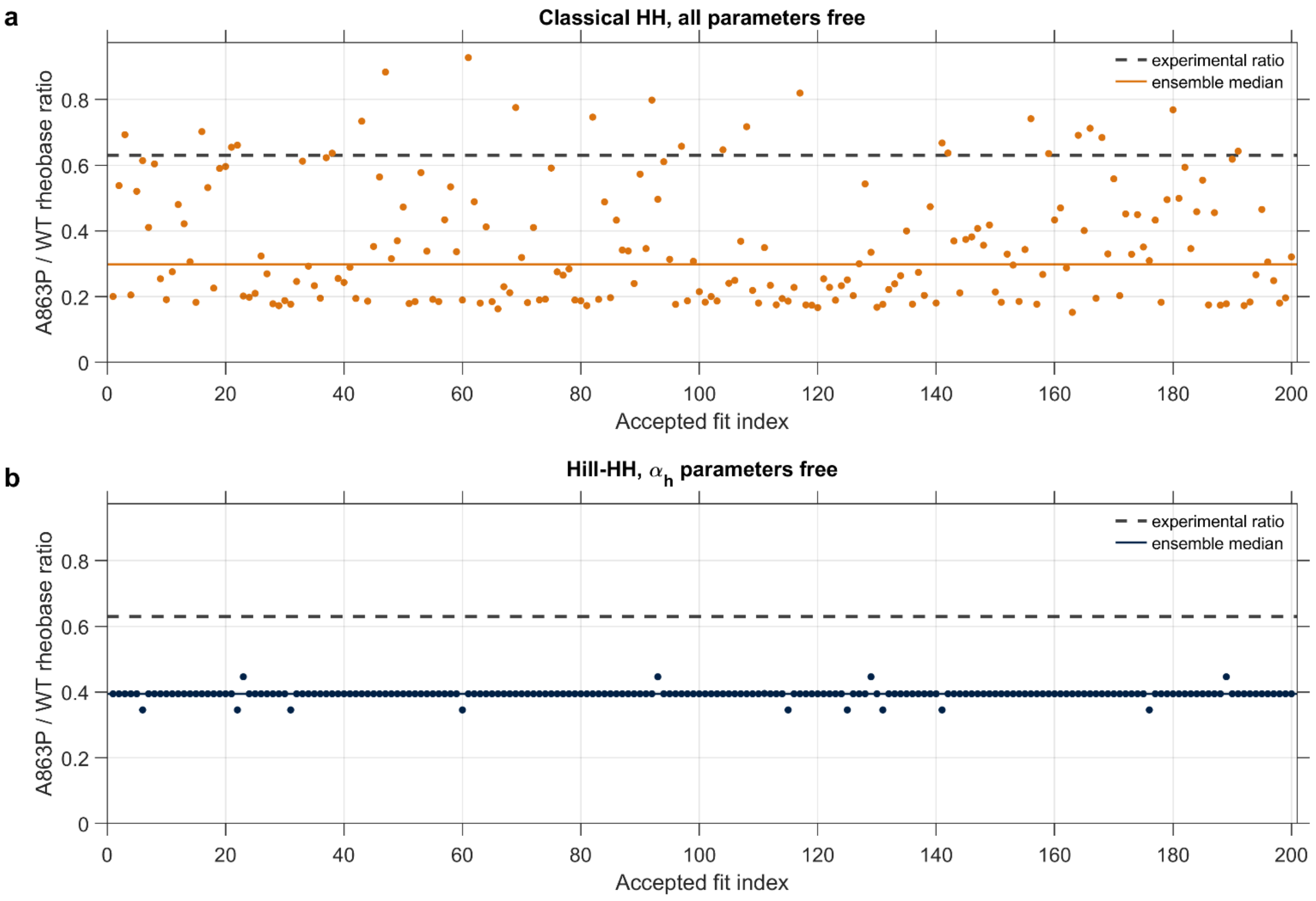


**Supplementary Figure 6. Variability in A863P rheobase predictions across accepted fitting ensembles.** Mutant-to-wild-type (WT) rheobase ratios for 200 accepted fits in each implementation. **a**, Classical Hodgkin-Huxley (HH) ensemble with all sodium-gating rate parameters free. **b**, Hill-based HH ensemble with the amplitude and voltage scale of the complementary inactivation rate $\alpha_h$ free and the complementary activation rate $\beta_m$ retained at its classical form. Each point represents one accepted fit; the horizontal axis indexes fits and does not denote a parameter sweep. Solid colored lines indicate ensemble medians, and dashed gray lines mark the experimental A863P/WT current-threshold ratio (approximately 0.633). Classical HH ratios span approximately 0.152-0.927, with a median of 0.298; Hill-based ratios span 0.346-0.446, with a median of 0.395. Classical fits use published Boltzmann activation and inactivation targets, whereas Hill-based fits use a Boltzmann activation target and the inactivation curve selected in the direct Hill implementation. The narrower Hill-based distribution therefore reflects repeatability under its specified targets and rate assumptions, rather than uniquely determined kinetics or superior quantitative prediction. Implementation details are provided in Supplementary Table S7.

| Gate | $A$ | $B$ | $C$ | $D$ | $E$ |
|---|---|---|---|---|---|
| $m$ | 0.100 | 25 | 10 | 4.000 | 18 |
| $n$ | 0.010 | 10 | 10 | 0.125 | 80 |
| $h$ | 1.000 | 30 | 10 | 0.070 | 20 |

**Supplementary Table S1.** Parameters of the classical Hodgkin-Huxley rate functions. Parameter symbols correspond to the rate expressions defined in the Methods.

| Quantity | Symbol | Value | Units |
|---|---|---|---|
| Membrane capacitance | $C_m$ | 1.0 | $\mu F/cm^2$ |
| Maximum $Na^+$ conductance | $\bar{g}_{Na}$ | 120 | $mS/cm^2$ |
| Maximum $K^+$ conductance | $\bar{g}_K$ | 36 | $mS/cm^2$ |
| Leak conductance | $\bar{g}_L$ | 0.3 | $mS/cm^2$ |
| $Na^+$ reversal potential | $E_{Na}$ | 50 | mV |
| $K^+$ reversal potential | $E_K$ | −77 | mV |
| Leak reversal potential | $E_L$ | −54.4 | mV |

**Supplementary Table S2.** Baseline membrane and conductance parameters of the Hodgkin-Huxley model.

| $V$ (mV) | $n_\infty$ | $m_\infty$ | $\beta_h\ (ms^{-1})$ |
|---|---|---|---|
| 0 | (0.315) | (0.042) | — |
| 6 | 0.448 | 0.103 | (0.09) |
| 10 | 0.496 | 0.145 | (0.13) |
| 19 | 0.674 | 0.323 | (0.40) |
| 26 | 0.728 | 0.569 | (0.65) |
| 32 | 0.772 | 0.709 | 0.75 |
| 38 | 0.806 | 0.778 | 0.79 |
| 51 | 0.859 | 0.895 | 0.94 |
| 63 | 0.891 | 0.963 | 1.19 |
| 76 | 0.915 | 0.975 | 1.19 |
| 88 | 0.935 | 1.029 | 1.50 |
| 100 | 0.953 | 0.997 | 1.50 |
| 109 | 0.961 | 0.980 | 1.50 |
| (∞) | (1.000) | (1.000) | — |

**Supplementary Table S3.** Historical Hodgkin-Huxley gating data at discrete transformed voltages used for sigmoidal fitting. All values are taken unchanged from Tables 1 and 2 of Hodgkin and Huxley (1952), with only the sign of V reversed to match the sign convention adopted throughout this work. Parentheses are kept as in the source: they mark limiting values as V tends to $\infty$ and values that Hodgkin and Huxley did not treat as reliable independent measurements.

| Variable | Model | SSE | AD p-value | AIC |
|---|---|---|---|---|
| $n_\infty$ | Boltzmann | 0.012396 | 0.0178* | −78.504 |
| | tanh | 0.012396 | 0.0177* | −78.504 |
| | arctan | 0.001816 | 0.3364 | −99.555 |
| | erf | 0.018572 | 0.1021 | −73.652 |
| | Hill | 0.001296 | 0.1952 | −103.600 |
| $m_\infty$ | Boltzmann | 0.008200 | 0.0374* | −91.791 |
| | tanh | 0.008200 | 0.0415* | −91.791 |
| | arctan | 0.007386 | 0.3579 | −91.150 |
| | erf | 0.010491 | 0.1352 | −88.588 |
| | Hill | 0.004858 | 0.2392 | −96.597 |
| $\beta_h$ | Boltzmann | 0.035450 | 0.7411 | −32.999 |
| | tanh | 0.035798 | 0.6186 | −32.930 |
| | arctan | 0.039773 | 0.9117 | −30.193 |
| | erf | 0.040453 | 0.8535 | −32.075 |
| | Hill | 0.031594 | 0.8105 | −31.805 |

**Supplementary Table S4.** Comparison of sigmoidal fits using the sum of squared errors (SSE), Anderson-Darling (AD) test, and Akaike information criterion (AIC). Lower SSE and AIC values indicate better fit and greater relative support, respectively, for fits to the same response data. An AD p-value below 0.05 indicates rejection of the tested distributional assumption; a value above 0.05 does not establish that assumption. Asterisks identify p-values below 0.05.

| Parameter | Minimum | Maximum |
|---|---|---|
| $a_m$ | −160.0 | −70.5 |
| $a_n$ | −66.0 | 2.0 |
| $a_h$ | −184.0 | 28.0 |
| $\ell_m$ | 6.5 | 12.5 |
| $\ell_n$ | 2.0 | 4.2 |
| $\ell_h$ | 9.0 | 129.0 |
| $V_{half,m}$ | 11.5 | 26.5 |
| $V_{half,n}$ | 6.5 | 38.0 |
| $V_{half,h}$ | 18.5 | 70.5 |

**Supplementary Table S5.** Hill-based model parameter ranges used for the spike-feature sensitivity analysis. These model-specific ranges were selected to retain non-damping trains with at least three spikes above 0 mV.

| Parameter | Minimum | Maximum |
|---|---|---|
| $A_m$ | 0.080 | 0.460 |
| $B_m$ | −8.0 | 29.0 |
| $C_m$ | 9.0 | 28.5 |
| $D_m$ | 1.5 | 5.3 |
| $E_m$ | 9.5 | 32.5 |
| $A_n$ | 0.003 | 0.012 |
| $B_n$ | 5.5 | 72.0 |
| $C_n$ | 0.0 | 12.0 |
| $D_n$ | 0.100 | 0.380 |
| $E_n$ | 20.0 | 100.0 |
| $A_h$ | 0.400 | 2.100 |
| $B_h$ | 18.5 | 67.0 |
| $C_h$ | 0.0 | 20.5 |
| $D_h$ | 0.030 | 0.580 |
| $E_h$ | 1.0 | 100.0 |

**Supplementary Table S6.** Classical Hodgkin–Huxley rate-parameter ranges used for the spike-feature sensitivity analysis. These model-specific ranges were selected to retain non-damping trains with at least three spikes above 0 mV.

| Model / implementation | Input data used | Extra assumptions required | WT rheobase ($\mu A/cm^2$) | A863P rheobase ($\mu A/cm^2$) | A863P/WT rheobase ratio | Spikes at $I_{app}$ = 40 (WT / A863P) | Interpretation |
|---|---|---|---|---|---|---|---|
| Hill-HH, single run (direct mapping) | Published $V_{1/2}$ and k of activation; published $V_{1/2}$ of inactivation (WT, A863P) | Classical $\beta_m$ and $\alpha_h$ retained; $\beta_h$ Hill parameters chosen so the $h_\infty$ midpoint equals the published $V_{1/2}$; no fitting | 17.69 | 7.19 | 0.406 | 1 / 16 (A863P 160 Hz) | Single prediction conditional on the specified mapping and kinetic assumptions. Expected direction of the threshold change reproduced. Firing reflects the expected phenotype: repetitive firing in A863P against a single spike in WT |
| Hill-HH ensemble, complementary rates fixed | Boltzmann $m_\infty$ (published $V_{1/2}$, k); chosen $h_\infty$ of the single run (published $V_{1/2}$) | $\beta_m$ and $\alpha_h$ classical; Hill parameters of $m_\infty$ and $\beta_h$ refitted; 200 random starts (200 accepted) | 17.50 [17.50–17.50] | 6.91 [6.91–6.91] | 0.395 [0.395–0.395] | 1 / 16 in all 200 fits | All 200 accepted fits give the same reported prediction across the tested starting points. Expected direction reproduced in every fit; firing reflects the expected phenotype (1 / 16 throughout) |
| Hill-HH ensemble, $\alpha_h$ parameters free | Boltzmann $m_\infty$ (published $V_{1/2}$, k); chosen $h_\infty$ of the single run (published $V_{1/2}$) | $\beta_m$ classical ($m_\infty$ fit does not constrain $\beta_m$, $\beta_m$ was retained); $\alpha_h$ amplitude and voltage scale free with $\beta_h$ Hill parameters; 200 random starts (200 accepted) | 17.50 [15.50–17.50] | 6.91 [6.05–6.92] | 0.395 [0.346–0.446] | 1 / 15–16 in 196 of 200 fits (A863P 150–160 Hz) | Expected direction reproduced in every fit, with the ratio varying slightly. Firing reflects the expected phenotype: repetitive firing in A863P in every fit |
| Classical HH, all parameters free | Published $V_{1/2}$ and k of $m_\infty$ and $h_\infty$ (WT, A863P) | All five parameters of $\alpha$ and $\beta$ (m, h) fitted to $x_\infty$ only, no kinetic data; 200 random starts | 45.20 [35.80–46.70] | 13.30 [6.94–41.90] | 0.298 [0.152–0.927] | 0 / 1 in 191 of 200 fits (WT 0–1, A863P 0–3) | Expected direction reproduced in all 200 fits, but the magnitude varies across a wide range with fits that reproduce the same $x_\infty$. Firing does not reflect the expected phenotype: no fit produces repetitive firing in A863P |
| Classical HH, complementary rates fixed | Published $V_{1/2}$ and k of $m_\infty$ and $h_\infty$ (WT, A863P) | $\beta_m$ and $\alpha_h$ fixed at classical values; other rate fitted to $x_\infty$ only; 200 random starts (104 accepted) | 47.20 [47.20–47.20] | 47.00 [47.00–47.00] | 0.996 [0.996–0.996] | 0 / 0 in all 104 fits | Identical in all 104 accepted fits, with a negligible threshold reduction (ratio ≈ 1). Both conditions are silent at the tested current. |

| Model / implementation | Input data used | Extra assumptions required | WT rheobase ($\mu A/cm^2$) | A863P rheobase ($\mu A/cm^2$) | A863P/WT rheobase ratio | Spikes at $I_{app}$ = 40 (WT / A863P) | Interpretation |
|---|---|---|---|---|---|---|---|
| Classical HH, voltage shift only | Published $V_{1/2}$ of $m_\infty$ and $h_\infty$ (WT, A863P); slope not used | Classical rates shifted along V to the published $V_{1/2}$; classical curve shapes and time-constant profiles retained under the corresponding voltage shifts; $n_\infty$ shifted by the WT $m_\infty$ shift (−0.88 mV) | 37.75 | 12.41 | 0.329 | 1 / 1 | Expected direction reproduced. Firing does not reflect the expected phenotype: a single spike in both conditions |
| Classical HH, tau-constrained | Published $V_{1/2}$ and k of $m_\infty$ and $h_\infty$ (WT, A863P) | $\tau(V)$ kept near classical HH $\tau$ shifted to the published $V_{1/2}$; constraint weight $\lambda$ (0.02–0.1); classical $K^+$ gate | 39.16 (37.47–42.31) | 16.16 (13.91–22.66) | 0.413 (0.370–0.535) | 1 / 1 ($\lambda$ = 0.02: 0 / 1) | Expected direction reproduced, but the magnitude depends on $\lambda$, which the data do not specify. Firing does not reflect the expected phenotype: neither condition fires repetitively |
| Classical HH, tau-rescaling | Published $V_{1/2}$ and k of $m_\infty$ and $h_\infty$ (WT, A863P) | Base kinetics from the tau-constrained fit ($\lambda$ = 0.05); $\alpha$ and $\beta$ of m and h scaled by $\varphi_m$, $\varphi_h \in$ {0.25, 0.5, 1, 2, 4} (25 combinations) | [19.30–47.10] | [5.94–46.70] | Median 0.460 Min 0.214 Max 0.991 ($\varphi$ = 1: 0.413) | 0–1 / 0–3 | Identical $x_\infty$ in all 25 cases; direction preserved (25/25), but magnitude varies across a wide range with kinetics not specified by the selected steady-state inputs |
| Classical HH fitted to Hill-derived $h_\infty$ (consistency check) | Published $V_{1/2}$ and k of $m_\infty$; chosen $h_\infty$ in the Hill-HH model | Complementary rates fixed; $h_\infty$ target is a model output, not published data | 17.00 | 6.75 | 0.397 | 1 / 15 | Matches Hill-HH by construction; not an independent biological prediction. Expected direction reproduced, as expected from a target taken from the Hill-HH model |

**Supplementary Table S7.** A863P implementation and kinetic-sensitivity summary. Rows specify the input targets, additional modeling assumptions, wild-type (WT) and mutant rheobases, mutant/WT ratios, and spike counts at $I_{app}$ = 40 $\mu A/cm^2$. Spike counts refer to the 100-ms current step. The experimental mutant/WT current-threshold ratio is approximately 0.63. Hill ensembles use the selected direct Hill inactivation curve, whereas most classical fits use published Boltzmann inactivation targets; these ensembles therefore do not constitute a comparison with identical targets. The fit to Hill-derived inactivation is retained as a consistency check. Each rheobase and ratio cell gives a central estimate followed by a bracketed range, but the meaning depends on the row. For the random-start ensemble rows the central value is the median over accepted fits and the bracket is the minimum to maximum across them; wild-type and mutant are fitted within the same start and every ratio is formed within a single paired fit, so the ratio range is a range over paired predictions and not a ratio of the separate wild-type and mutant ranges. For the time-constant-constrained row the reference value is the reference fit at λ = 0.05 and the parenthesized range spans the three constraint weights $\lambda$ = 0.02, 0.05 and 0.1; it is not an ensemble median. For the time-constant-rescaling row the ratio column reports the median over the 25 ($\emptyset_m, \emptyset_h$) combinations, the reported minimum and maximum span these combinations, and the value in parentheses is the unscaled reference at $\emptyset_m = \emptyset_h$= 1, which coincides with the $\lambda$ = 0.05 fit by construction. Rows describing a single deterministic implementation report single values without ensemble ranges. Ensemble sizes and acceptance counts are listed where applicable.

| Mutation | Condition | Gate | Published $V_{1/2}$ (mV) | Published k (mV) | Source of k | Hill a (mV) | Hill $V_{half}$(mV) | Hill $\ell$ |
|---|---|---|---|---|---|---|---|---|
| A863P | WT | Activation | −15.9 ± 0.8 | 7.3 ± 0.1 | Reported | −250 | −15.9 | 32.0685 |
| A863P | A863P | Activation | −23.6 ± 0.6 | 8.4 ± 0.2 | Reported | −250 | −23.6 | 26.9524 |
| A863P | WT | Inactivation | −76.9 ± 0.8 | 6.9 ± 0.1 | Reported | −150 | −73 | 4.88 |
| A863P | A863P | Inactivation | −67.1 ± 0.9 | 6.4 ± 0.2 | Reported | −150 | −47 | 4.68 |
| F1449V | WT | Activation | −15.2 ± 1.3 | 7.65 | Recovered | −250 | −15.2 | 30.6760 |
| F1449V | F1449V | Activation | −22.8 ± 1.3 | 8.08 | Recovered | −250 | −22.8 | 28.1319 |
| F1449V | WT | Inactivation | −71.3 ± 0.8 | 6.68 | Recovered | −150 | −57.5 | 4.4 |
| F1449V | F1449V | Inactivation | −67.0 ± 1.4 | 7.00 | Recovered | −150 | −39 | 3.5 |
| A1632G | WT | Activation | −20.68 | 8.09 | Recovered | −250 | −20.68 | 28.3461 |
| A1632G | A1632G | Activation | −27.17 | 8.64 | Recovered | −250 | −27.17 | 25.7905 |
| A1632G | WT | Inactivation | −86.32 | 7.58 | Recovered | −130 | −96.576 | 3.4 |
| A1632G | A1632G | Inactivation | −76.91 | 6.75 | Recovered | −120 | −70.0242 | 2 |

Supplementary Table S8. Published gating descriptors (Dib-Hajj et al. 2005; Harty et al. 2006; Yang et al. 2016) and selected Hill parameters for A863P, F1449V, and A1632G, with study-specific wild-type (WT) controls. Published values are reproduced as printed, and the $\pm$ values are the uncertainties reported in the source studies. Half-activation and half-inactivation voltages are reported in all three studies and were used as reported. Among the descriptors used here, slope factors are directly reported for A863P; those marked “recovered” were estimated by reading the normalized value of the published curve at a voltage away from the midpoint and solving the Boltzmann equation for the slope factor. Activation descriptors are mapped under the specified Hill shift parameter a; inactivation parameters depend on the chosen rate construction. *The Hill parameters for inactivation describe $\beta_h$, not $h_\infty$ itself.*

| Mutation | Condition | Published $V_{1/2}$ (mV) | Achieved $V_{1/2}$ (mV) | Error (mV) |
|---|---|---|---|---|
| A863P | WT | −76.90 | −76.74 | +0.16 |
| A863P | A863P | −67.10 | −66.87 | +0.23 |
| F1449V | WT | −71.30 | −71.11 | +0.19 |
| F1449V | F1449V | −67.00 | −66.79 | +0.21 |
| A1632G | WT | −86.32 | −86.39 | −0.07 |
| A1632G | A1632G | −76.91 | −76.40 | +0.51 |

**Supplementary Table S9.** Published and achieved half-inactivation voltages of the $h_\infty$ derived by the modified model. Inactivation is represented by a Hill function for $\beta_h$ together with the classical $\alpha_h$, so that $h_\infty = \alpha_h/(\alpha_h + \beta_h)$. The parameters of $\beta_h$ were chosen so that the midpoint of the resulting $h_\infty$ matches the published value for each condition. *The* resulting midpoints differ from *the* published values by no more than *0.51* mV. Error is the achieved minus the published midpoint; agreement *in* midpoint alone does not establish agreement in slope or kinetics.

| Case study | Hill a (mV) | Hill $\ell$ | Hill $V_{1/2}$ (mV) | Equivalent k (mV) |
|---|---|---|---|---|
| A863P | −110 | 6.333 | −15 | 15.00 |
| F1449V | −110 | 6.333 | −15 | 15.00 |
| A1632G | −130 | 7 | −21 | 15.57 |

**Supplementary Table S10.** The potassium gate of the modified model. Potassium-gating parameters were specified by design rather than fitted to mutation-specific potassium measurements, and were held identical between wild type and mutant within each case study.

| Mutation | Perturbation | WT rheobase ($\mu A/cm^2$) | Mutant rheobase ($\mu A/cm^2$) | Mutant / WT ratio |
|---|---|---|---|---|
| A863P | Activation shift only | 17.69 | 8.12 | 0.459 |
| A863P | Inactivation shift only | 17.69 | 16.25 | 0.919 |
| A863P | Both shifts combined | 17.69 | 7.19 | 0.406 |
| F1449V | Activation shift only | 17.31 | 8.75 | 0.505 |
| F1449V | Inactivation shift only | 17.31 | 15.81 | 0.913 |
| F1449V | Both shifts combined | 17.31 | 7.97 | 0.460 |
| A1632G | Activation shift only | 13.78 | 5.84 | 0.424 |
| A1632G | Inactivation shift only | 13.78 | 12.41 | 0.900 |
| A1632G | Both shifts combined | 13.78 | 5.00 | 0.363 |

**Supplementary Table S11.** Direct Hill-based rheobase predictions for activation-only, inactivation-only, and combined perturbations in A863P, F1449V, and A1632G. *Each ratio is the mutant rheobase divided by the* corresponding wild-type (WT) rheobase, rounded *to three* decimal places. Activation perturbations account *for most of the predicted* threshold reduction; inactivation perturbations alone reduce rheobase by approximately 8-10%. Perturbations use the descriptors and parameter choices *in* Supplementary Table S8.

| Mutation | Experimental ratio | Experimental values | Source | Predicted ratio (both shifts) |
|---|---|---|---|---|
| A863P | 0.633 | 157 / 248 pA | Harty et al., J Neurosci 2006 | 0.406 |
| F1449V | 0.750 | 93.1 / 124.1 pA | Dib-Hajj et al., Brain 2005;128:1847–1854 | 0.460 |
| A1632G | 0.646 | 102 / 158 pA | Yang et al., J Neurosci 2016;36(28):7511–7522 | 0.363 |

**Supplementary Table S12.** Experimental current-threshold *ratios* and direct Hill-based rheobase ratios for A863P, F1449V, and A1632G. Ratios are mutant divided by the study-specific wild-type (WT) value; model predictions include both activation and inactivation perturbations. Predicted ratios (0.363-0.460) are lower than experimental ratios (0.633-0.750), indicating *an* overestimated threshold *reduction in* all three benchmarks. Experimental currents and model current densities are compared through dimensionless ratios. These results do not establish *the* cause of *the* quantitative discrepancy.

| Mutation | Condition | Threshold (mV) | Amplitude (mV) | AHP (mV) | Half-width (ms) |
|---|---|---|---|---|---|
| F1449V | WT | −54.40 | 95.90 | no AHP | 1.60 |
| F1449V | F1449V | −54.38 | 102.05 | 6.19 | 1.70 |
| A1632G | WT | −54.43 | 90.07 | 0.63 | 1.30 |
| A1632G | A1632G | −54.29 | 98.89 | 6.29 | 1.30 |

**Supplementary Table S13.** Simulated first-spike features of F1449V and A1632G and their respective wild-type (WT) controls at $I_{app}$ = 40 $\mu A/cm^2$. *Features are measured from the first spike* during *the 100-ms* current *step and* reported *to two* decimal places. AHP denotes afterhyperpolarization. Voltage thresholds change little, while spike amplitudes increase by approximately 6% for F1449V *and* 10% *for* A1632G. Half-width changes from 1.60 to 1.70 ms for F1449V *and* remains 1.30 ms for A1632G; AHP increases in both mutants. These deterministic comparisons do *not* establish statistical equivalence to experimental recordings.

| Gate | Parameter | Role | Lower bound | Upper bound |
|---|---|---|---|---|
| Activation | a | Hill offset voltage of $m_\infty$ | −400 | −250 |
| Activation | $\ell$ | Hill coefficient of $m_\infty$ | 2 | 80 |
| Activation | V_half | half-activation voltage of $m_\infty$ | −50 | 10 |
| Inactivation | a | Hill offset voltage of $\beta_h$ | −300 | −150 |
| Inactivation | $\ell$ | Hill coefficient of $\beta_h$ | 1 | 60 |
| Inactivation | V_half | half-point voltage of $\beta_h$ | −110 | 10 |
| Inactivation | $D_h$ | amplitude of the classical $\alpha_h$ | 0.014 | 0.35 |
| Inactivation | $E_h$ | voltage scale of the classical $\alpha_h$ | 4 | 100 |

**Supplementary Table S14.** Parameter bounds of the random-start ensemble of the modified model. Bounds are fixed across all draws. Starting points were drawn independently within these bounds for every fit, log-uniformly for the strictly positive parameters and uniformly for the voltage parameters.

| Gate | Parameter | Role in the rate expression | Lower bound | Upper bound |
|---|---|---|---|---|
| Activation | $A_m$ | amplitude of $\alpha_m$ | 0.001 | 50 |
| Activation | $B_m$ | voltage offset of $\alpha_m$ | −200 | 300 |
| Activation | $C_m$ | voltage scale of $\alpha_m$ | 1 | 60 |
| Activation | $D_m$ | amplitude of $\beta_m$ | 0.001 | 100 |
| Activation | $E_m$ | voltage scale of $\beta_m$ | 2 | 200 |
| Inactivation | $D_h$ | amplitude of $\alpha_h$ | 0.00001 | 10 |
| Inactivation | $E_h$ | voltage scale of $\alpha_h$ | 2 | 200 |
| Inactivation | $B_h$ | voltage offset of $\beta_h$ | −200 | 300 |
| Inactivation | $C_h$ | voltage scale of $\beta_h$ | 1 | 60 |
| Inactivation | $A_h$ | amplitude of $\beta_h$ | 0.001 | 50 |

**Supplementary Table S15.** Parameter bounds of the time-constant-constrained classical fits. These are fixed absolute intervals, identical for every gate, condition and value of the constraint weight, and differ from the bounds of the random-start ensembles. *Rate parameters are evaluated at V + 40 mV.*
